# Why Ortho- and Para-Hydroxy Metabolites Can Scavenge Free Radicals That the Parent Atorvastatin Cannot? Important Pharmacologic Insight from Quantum Chemistry

**DOI:** 10.1101/2022.07.29.502012

**Authors:** Ioan Bâldea

**Affiliations:** Theoretical Chemistry, Heidelberg University, Im Neuenheimer Feld 229, D-69120 Heidelberg, Germany

**Keywords:** Radical-scavenging activity, atorvastatin, antioxidant mechanisms, HAT, SPLET, SET-PT, global chemical reactivity indices, DPPH radical, solvent effects, quantum chemistry

## Abstract

The pharmaceutical success of atorvastatin (ATV), a widely employed drug against the “bad” cholesterol (LDL) and cardiovascular diseases, traces back to its ability to scavenge free radicals. Unfortunately, information on its antioxidant properties is missing or unreliable. Here, we report detailed quantum chemical results for ATV and its ortho- and para-hydroxy metabolites (o-ATV, p-ATV) in methanolic phase. They comprise global reactivity indices bond order indices and spin densities as well as all relevant enthalpies of reaction (bond dissociation BDE, ionization IP and electron attachment EA, proton detachment PDE and proton affinity PA, and electron transfer ETE). With these properties in hand, we can provide the first theoretical explanation of the experimental finding that, due to their free radical scavenging activity, ATV hydroxy metabolites rather than the parent ATV have substantial inhibitory effect on LDL and the like. Surprisingly (because it is contrary to the most cases currently known), we unambiguously found that HAT (direct hydrogen atom transfer) rather than SPLET (sequential proton loss electron transfer) or SET-PT (stepwise electron transfer proton transfer) is the thermodynamically preferred pathway by which o-ATV and p-ATV in methanolic phase can scavenge DPPH• (1,1-diphenyl-2-picrylhydrazyl) radicals.

## 1. Introduction

The highly radical scavenging active cholesterol-lowering drug atorvastatin (ATV) [1] is an outstanding success sale story [2]. It was patented in 1985 and approved by the Food and Drug Administration (FDA) in 1996 for medical use. Sold under the name of lipitor, it received record high revenues of about 12.8 billion US dollars in 2006, still generated 10 billion US dollars in the year of patent loss (2011) and nearly two billion US dollars in 2019. ATV, one of the most prescribed drugs in the US today, is mainly employed to prevent high risk for developing cardiovascular diseases and as treatment for abnormal lipid levels (dyslipidemia). ATV’s inhibition of the HMG-CoA (3 hydroxy-3-methylglutaryl coenzyme A) reductase is plausibly related to the high radical scavenging potency against lipoprotein oxidation.

ATV made the object of several theoretical investigations in the past [3,4]. Still, the antioxidant properties of ATV were only recently investigated from the quantum chemical perspective [5]. Unfortunately, as we drew attention recently [6], the only quantum chemical attempt of which we are aware [5] is plagued by severe flaws [6], and this makes mandatory the effort (undertaken in the present paper) of properly reconsidering the antioxidant capacity of ATV and its ortho- and para-hydroxy metabolites in methanol. For the notoriously poor soluble ATV, this solvent is of special interest. ATV is freely soluble in methanol. In addition, antioxidant assays are mostly done in methanolic environment [5,7]. Along with quantities traditionally related to the antioxidant activity, the present study will also reports on the ATV global chemical reactivity indices, relevant bond data as well as spin densities of radical species generated by H-atom abstraction from ATV and related ortho- and para-hydroxylated derivatives (o-ATV, p-ATV, respectively).

Theoretical understanding of the differences between ATV and its ortho- and parahydroxy metabolites, which is missing to date, is of paramount practical importance. A twenty four years old experimental study reported that atorvastatin ortho- and parahydroxy metabolites (o-ATV and p-ATV, respectively) protect, e.g., LDL from oxidation, while the parent ATV does not [8]. Importantly for the results we are going to present in Section 3.5, the free radical scavenging activity of o-ATV and p-ATV was analyzed by the ubiquitous 1,1 diphenyl-2 picryl-hydrazyl (DPPH•) assay in ref. [8]. Our study is able to provide the first theoretical explanation of this experimental finding.

## 2. Computational details

The results reported below were obtained from quantum chemical calculations wherein all necessary steps (geometry optimizations, frequency calculations, and electronic energies) where conducted at the same DFT level of theory by running GAUSSIAN 16 [9] on the bwHPC platform [10]. In all cases investigated, we convinced ourselves that all frequencies are real. In all calculations we used 6-31+G(d,p) basis sets [11,12] and, unless otherwise specified (see Tables 2 and 5), the hybrid B3LYP exchange correlation functional [13–16].

For comparative purposes, we also present results obtained by using the PBE0 [17] functional (Tables 2 and 5) and Truhlar’s M062x [18,19] (Table 2). Computations for open shell species were carried out using unrestricted spin methods (e.g., UB3LYP and UPBE0). In most radicals, employing the more computationally demanding quadratic convergence SCF methods was unavoidable. We convinced ourselves that spin contamination is not a severe issue. In all these calculations, we invariably found a value ⟨ *S*^2^⟩= 0.7501 for the total spin after annihilation of the first spin contaminant, versus the exact value ⟨*S*^2^⟩ = 3/4.

Still, to better check this aspect, for ATV’s cation and anion as well as for the ATV1H and ATV4H radicals (see Section 3.1 for the meaning of these acronyms) we also undertook the rare numerical effort (enormous for molecules with almost 80 atoms) of performing *full* restricted open shell (ROB3LYP) calculations; that is, not only single point calculations for electronic energy but also geometry optimization and (numerical) vibrational frequency calculations, and all these in solvent. Differences between UB3LYP and ROB3LYP were reasonably small (see Tables 2 and 5), but they should make it clear that claims (often formulated in the literature on antioxidation) of chemical accuracy (∼ 1 kcal/mol) at the B3LYP/6-31+G(d,p) are totally out of place. From experience with much smaller molecules and much simpler chemical structures (e.g. ref. [20]) we had to learn that achieving this accuracy for bond dissociation enthalpies and proton affinity (BDE and PA, quantities entering the discussion that follows) is often illusory even for extremely computationally demanding state-of-the-art compound model chemistries (CBS-QB3, CBS-APNO, G4, W1BD); see, e.g., Figure 10 of ref. [20]. DFT-calculations done by us and by others [21] revealed that, e.g., errors in ionization potential can amount up to 0.7 eV (16 kcal/mol) even when employing the functional B3LYP and the largest Pople basis set 6-311++G(3df,3pd).

Unless otherwise specified, the solvent (methanol) was accounted for within the polarized continuum model (PCM) [22] using the integral equation formalism (IEF) [23]. Although this is the “gold standard” for modeling solvents in the literature on free radical scavenging, one should be aware that this framework ignores specific solvation effects (hydrogen bonds). Because they may play an important role, e.g., in proton transfer reactions, theoretical estimates of PA may not be sufficiently accurate. While this makes comparison with experiment problematic, it should be a less critical issue when comparing among themselves PA values of various antioxidants in a given solvent (e.g., methanol). To better emphasize why we believe that solvent effects in the context of antioxidants deserve a more careful consideration, along with IEFPCM-based results, in Tables 2 and 5 we also present results obtained in Truhlar’s SMD solvation model [24–26].

GABEDIT [27] was used to generate Figures 1, 3, 4, and 5 from the GAUSSIAN output (*.log) files. To compute Wiberg bond order indices, we used the package NBO 6.0 [28] interfaced with GAUSSIAN 16. The reason why we use Wiberg bond order indices [29] rather than the heavily advertised Mayer bond order indices [30] was explained elsewhere [31]. All thermodynamic properties were calculated at *T* = 298.15 K.

**Figure 1.**
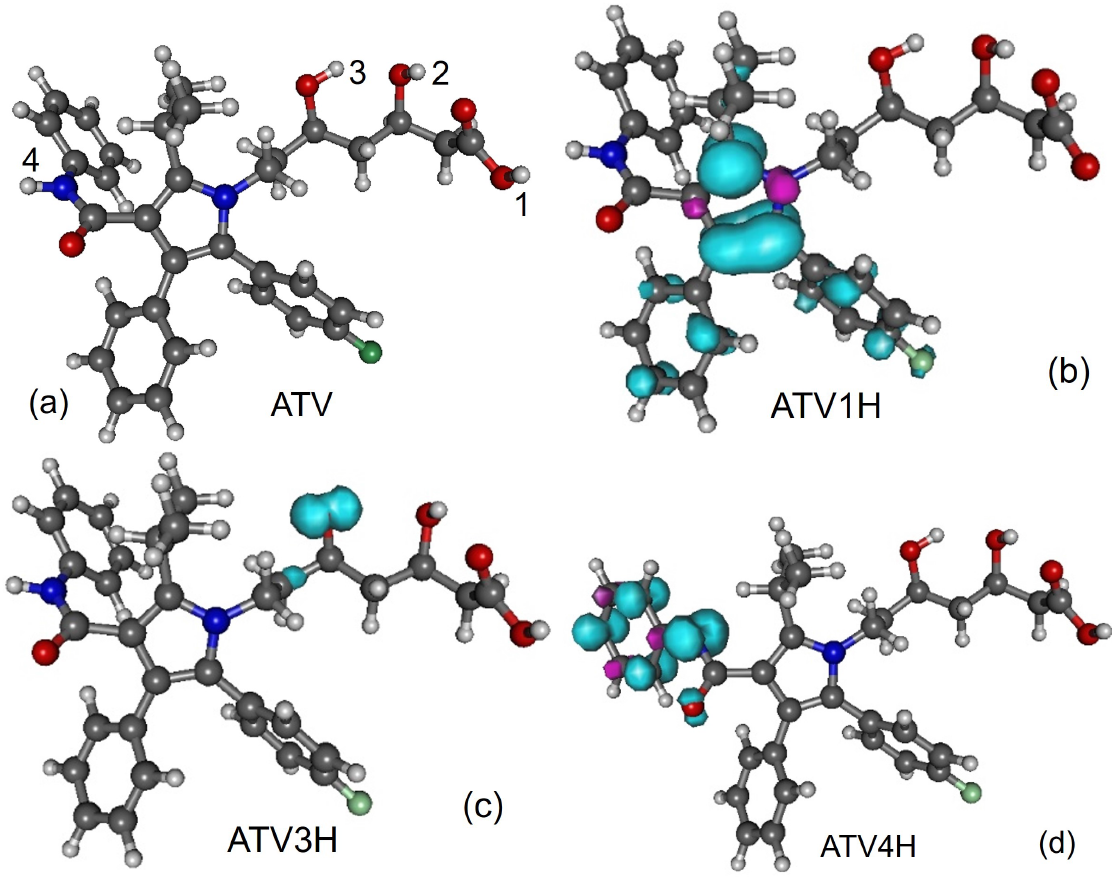
(a) Optimized ATV geometry. Spin densities of neutral radicals generated from it by H-atom abstraction at positions indicated in the inset: (b) ATV1H, (c) ATV3H, and (d) ATV4H.

**Figure 2.**
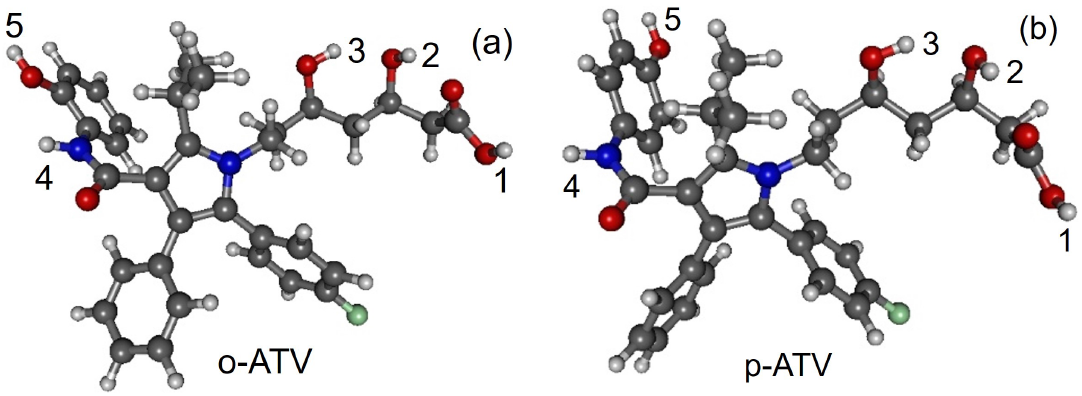
Optimized geometries of atorvastatin ortho- and para-hydroxy metabolites: (a) o-ATV and (b) p-ATV.

**Figure 3.**
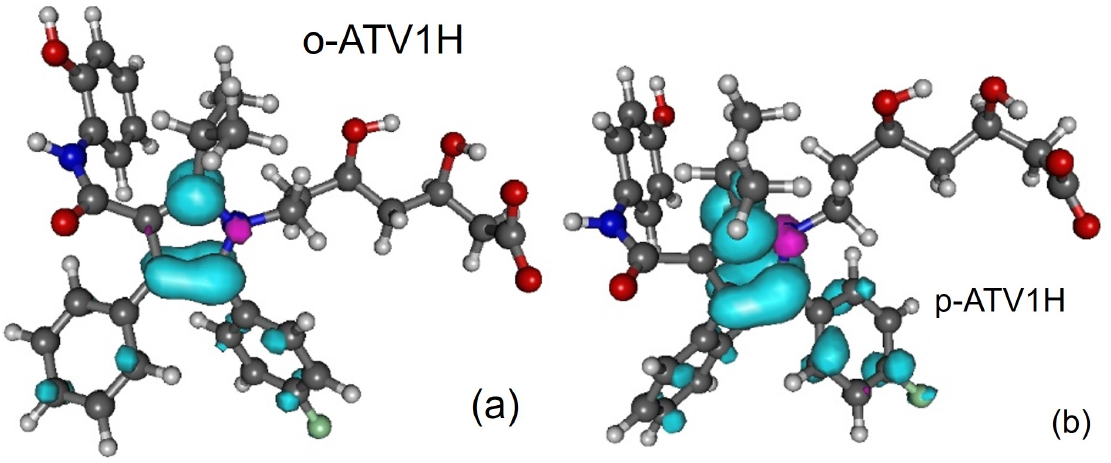
Spin densities of radicals generated by H-atom abstraction at position 1-OH: (a) o-ATV1H and (b) p-ATV1H.

**Figure 4.**
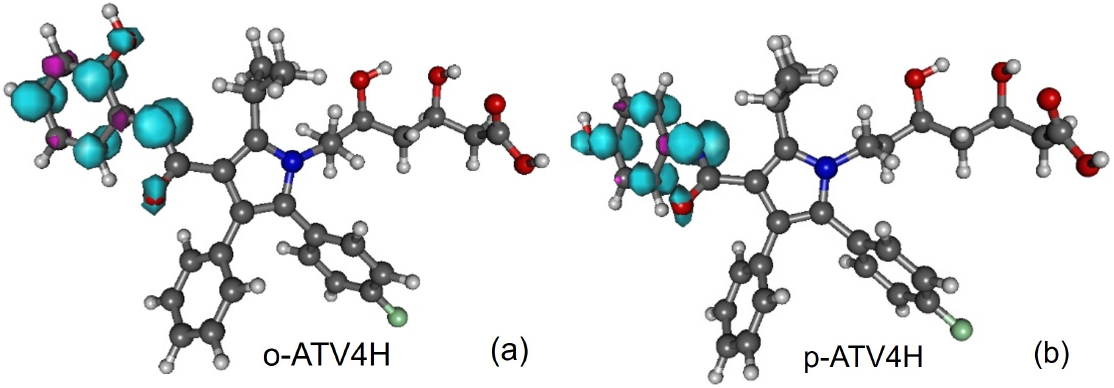
Spin densities of radicals generated from atorvastatin ortho- and para-hydroxy metabolites by H-atom abstraction at position 4-NH: (a) o-ATV4H and (b) p-ATV4H.

**Figure 5.**
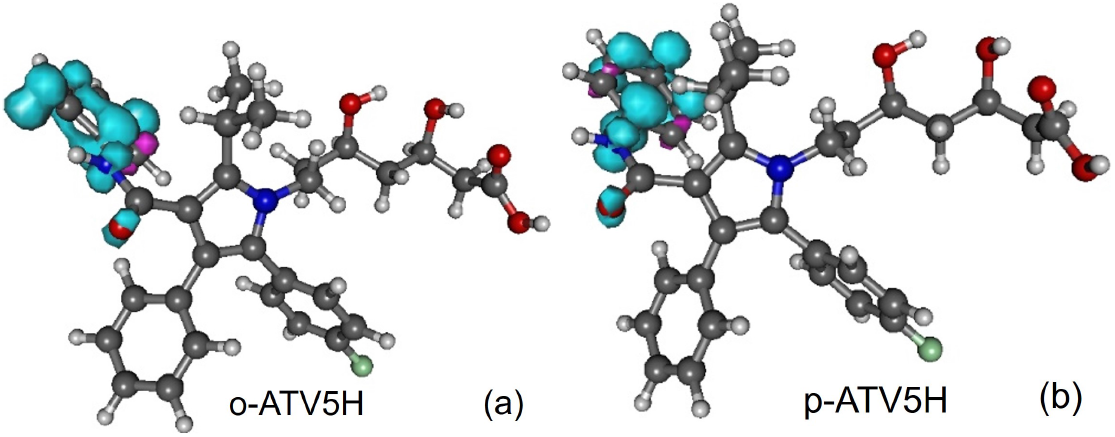
Spin densities of radicals generated from atorvastatin ortho- and para-hydroxy metabolites by H-atom abstraction at position 5-OH: (a) o-ATV5H and (b) p-ATV5H.

## 3. Results and Discussion

### 3.1. Molecular Geometries

Along with the neutral, cation and anion ATV — molecular formula C_33_H_35_FN_2_O_5_, IUPAC name (3R,5R)-7-[2-(4-fluorophenyl)-3-phenyl-4-(phenylcarbamoyl)-5-propan-2-yl-pyrrol-1-yl]-3,5-dihydroxyheptanoic acid, CAS number 134523-00-5— and its metabolites ortho-hydroxy atorvastatin (o-ATV and para–hydroxyatorvastatin (p-ATV), we also investigated the radicals (e.g., ATV*n*H•) generated by H-atom abstraction from their O – H and N – H groups as well as the anions ATV*n*H^−^ of the latter. Here, *n*(= 1, 2, 3, …) labels the various positions of the H-atoms, as depicted in Figures 1, 3, 4, and 5.

All quantities to be discussed below were calculated at the total electronic energy minima of the species listed above obtained via B3LYP/6-31+G(d,p)/IEFPCM optimization (cf. Section 2), which (with the grain of salt mentioned in the caption of Figure 7) posed no special problems. Neither H-atom abstraction nor ortho- and para-O – H addition spectacularly modifies the molecular conformation. *Z*-matrices for optimized geometries of representative species are presented in Tables A1, A2, A3 and A4 and Figures 1, 3, 4, and 5. Rather than Cartesian coordinates, we prefer to show *Z*-matrices because they facilitate comparison between various species and methods.

### 3.2. Chemical Reactivity Indices

The global chemical reactivity indices investigated in this work are listed below along with their expressions in terms of the ionization potential IP and electroaffinity EA [32–36]:

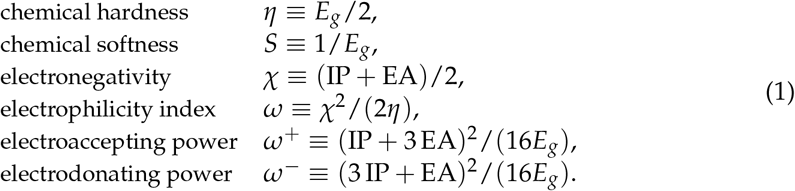

Here, *E*_*g*_ ≡ IP − EA is the fundamental (or transport) “HOMO-LUMO” gap [32,37,38]. Noteworthily, the presence of a solvent makes the popular approximation of IP and EA as the HOMO and LUMO energies with reversed sign (Koopmans theorem) totally inadequate [39]. Therefore, they were calculated in the usual way as enthalpy differences (cf. equations (3a) and (5)).

By and large, one may expect that these indices can give a flavor of the overall stability of a molecule and are useful in predicting how a certain chemical environment evolves in time [40,41]. In certain situations they turned out to be useful for comparing properties of different molecular species [33,42,43].

The presently calculated global chemical reactivity indices of ATV and its metabolites are collected in Table 1 and depicted in Figure 6. Having a chemical hardness *η* of about 2 eV, ATV, o-ATV, and p-ATV exhibit a good chemical stability. This value lies between the values of the natural antioxidants phenol and trolox, for which our calculations at the same B3LYP/6-31+G(d,p)/IEFPCM level yielded *η* = 2.56 eV and *η* = 1.88 eV, respectively. For all three species, the electrophilic indices [33,42,43] are *ω* ≈ 1.8 eV, a value exceeding the value of 1.50 eV, which is considered the threshold for strong electrophiles [44]. For comparison, let us again mention the values *ω* = 1.61 eV and *ω* = 1.85 eV, which we computed for phenol and trolox, respectively.

**Table 1.** Global chemical reactivity indices (eV) computed via B3LYP/6-31+G(d,p)/IEFPCM for atorvastatin and its ortho- and para-hydroxy metabolites and two natural oxidants in methanol.

| Molecule | IP | EA | $E_g$ | $\eta$ | $\mu$ | $S$ | $\omega$ | $\omega^+$ | $\omega^-$ |
| --- | --- | --- | --- | --- | --- | --- | --- | --- | --- |
| ATV | 4.64 | 0.72 | 3.92 | 1.96 | -2.68 | 0.26 | 1.83 | 0.74 | 3.42 |
| o-ATV | 4.64 | 0.73 | 3.90 | 1.95 | -2.68 | 0.26 | 1.85 | 0.75 | 3.43 |
| p-ATV | 4.60 | 0.67 | 3.93 | 1.97 | -2.64 | 0.25 | 1.77 | 0.70 | 3.34 |
| Phenol | 5.43 | 0.31 | 5.12 | 2.56 | -2.87 | 0.20 | 1.61 | 0.49 | 3.36 |
| Trolox | 4.51 | 0.76 | 3.76 | 1.88 | -2.63 | 0.27 | 1.85 | 0.77 | 3.40 |

**Table 2.** Global chemical reactivity indices (eV) for ATV in methanol computed using 6-31+G(d,p) basis sets and the exchange-correlation functionals (B3LYP, PBE0, M062x) and solvent models (IEFPCM, SMD) specified above.

| Molecule | Method | IP | EA | $E_g$ | $\eta$ | $\mu$ | $S$ | $\omega$ | $\omega^+$ | $\omega^-$ |
| --- | --- | --- | --- | --- | --- | --- | --- | --- | --- | --- |
| ATV | UB3LYP/IEFPCM | 4.64 | 0.72 | 3.92 | 1.96 | -2.68 | 0.26 | 1.83 | 0.74 | 3.42 |
|  | B3LYP/SMD | 4.39 | 0.63 | 3.76 | 1.88 | -2.51 | 0.27 | 1.67 | 0.65 | 3.16 |
|  | ROB3LYP/IEFPCM | 4.69 | 0.70 | 3.98 | 1.99 | -2.69 | 0.25 | 1.82 | 0.72 | 3.42 |
|  | UPBE0/IEFPCM | 4.67 | 0.71 | 3.96 | 1.98 | -2.69 | 0.25 | 1.82 | 0.73 | 3.41 |
|  | UM062x/IEFPCM | 4.95 | 0.73 | 4.22 | 2.11 | -2.84 | 0.24 | 1.91 | 0.75 | 3.59 |
| o-ATV | UB3LYP/IEFPCM | 4.64 | 0.73 | 3.90 | 1.95 | -2.68 | 0.26 | 1.85 | 0.75 | 3.43 |
|  | UB3LYP/SMD | 4.38 | 0.63 | 3.75 | 1.87 | -2.51 | 0.27 | 1.68 | 0.66 | 3.16 |
| p-ATV | UB3LYP/IEFPCM | 4.60 | 0.67 | 3.93 | 1.97 | -2.64 | 0.25 | 1.77 | 0.70 | 3.34 |
|  | UB3LYP/SMD | 4.37 | 0.61 | 3.77 | 1.88 | -2.49 | 0.27 | 1.65 | 0.64 | 3.13 |

**Figure 6.**
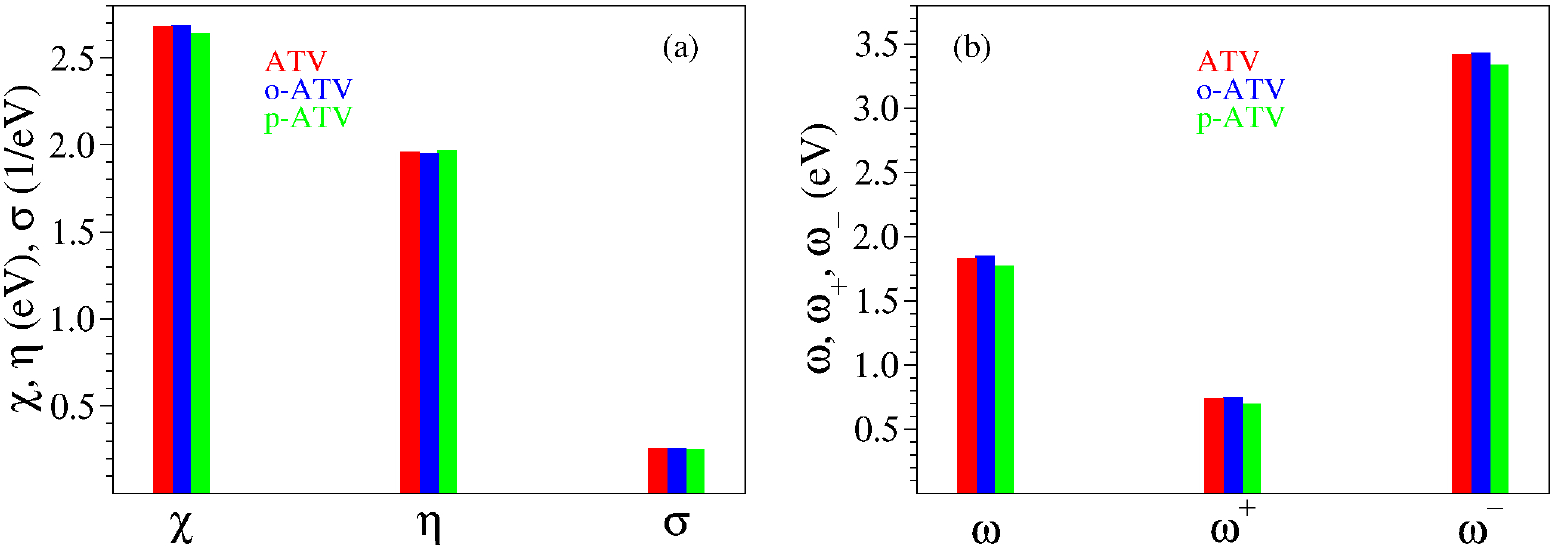
Global chemical reactivity indices defined by equation (1) for atorvastatin and its ortho- and para-hydroxy metabolites.

Inspection of Table 1 reveals that, similarly to the quantities *η* and *ω* considered above, all global chemical reactivity of ATV, o-ATV, and p-ATV are comparable to those of well known natural antioxidants. Could we then expect that ATV flavors (or other molecules) have indeed good antioxidant potency merely based on global chemical reactivity indices comparable to those of good antioxidants?

The analysis in the next section will unravel that, in fact, the global chemical indices have little relevance for assessing the antioxidant activity of a certain molecule. For the time being, let us remark that the values of Table 1 would rather suggest that ATV and o-ATV have similar antioxidant properties, and that ATV (possibly) performs (slightly) better than p-ATV.

### 3.3. Antioxidant Mechanisms and Pertaining Enthalpies of Reaction

As is widely discussed in the literature, an H-atom can be transferred to a free radical in one or two step processes. The three antioxidative mechanisms (HAT, SET-PT, and SPLET) and the corresponding reaction enthalpies (BDE, IP and PDE, PA and ETE, respectively) can be expressed as follows:

Direct hydrogen atom transfer (HAT) [45–47]

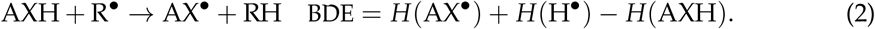

Stepwise electron transfer proton transfer (SET-PT) [48,49]

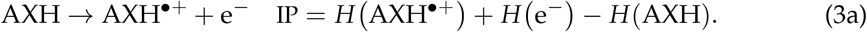

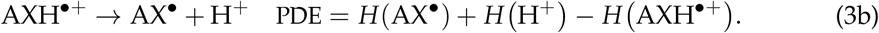

Sequential proton loss electron transfer (SPLET) [50,51]

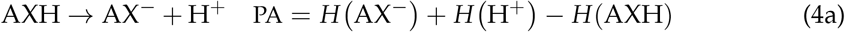

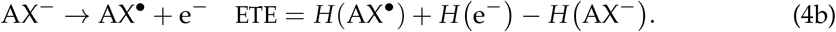

In our specific case, X stands for an O or an N atom.

Related to the above (albeit not directly entering the aforementioned antioxidation mechanisms), the electron attachment process is quantified by the electroaffinity defined as

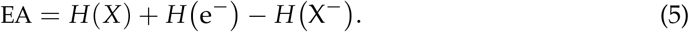

BDE, IP, PDE, PA, and ETE are enthalpies of reaction which can be obtained as adiabatic properties from standard Δ-DFT prescriptions [38,52,53]. To this aim, along with the enthalpies of the various ATV-based species entering the above reactions, the enthalpies of the H-atom, proton and electron in methanol are also needed [6]. They are presented in Table 3.

**Table 3.**
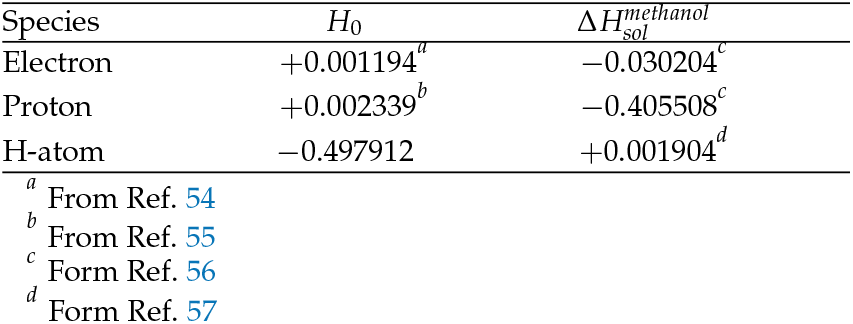
Gas phase enthalpies *H*_0_ and solvation enthalpies Δ*H*_*sol*_ in hartree utilized in the present calculations. For the for the gas phase enthalpy of the H-atom we used the value for the B3LYP/6-31+G(d,p) electronic energy (−0.500273 hartree) and the value of 1.4816 kcal/mol for thermal correction to enthalpy common for all compound model chemistries from GAUSSIAN 16.

| Species | $H_0$ | $\Delta H_{sol}^{methanol}$ |
| --- | --- | --- |
| Electron | +0.001194<sup>a</sup> | −0.030204<sup>c</sup> |
| Proton | +0.002339<sup>b</sup> | −0.405508<sup>c</sup> |
| H-atom | −0.497912 | +0.001904<sup>d</sup> |

<sup>a</sup> From Ref. 54<sup>b</sup> From Ref. 55<sup>c</sup> From Ref. 56<sup>d</sup> From Ref. 57

The presently computed thermodynamic parameters are collected in Table 4 and depicted in Figure 10. Inspection of Table 4 and Figure 10 reveals that the additional 5-OH group has no notable impact on the O-H bond cleavage at positions 1-OH, 2-OH, and 3-OH, neither homolytic and heterolytic. BDE for H-atom abstraction at positions 1-OH, 2-OH, and 3-OH in ATV, o-ATV and p-ATV is basically the same. The differences between the values calculated by us for ATV, o-ATV, and p-ATV amounting to at most 0.5 kcal/mol are certainly irrelevant; recall that we showed recently [20] that even for much smaller molecules in vacuo DFT/B3LYP calculations with the largest Pople basis set 6-311++G(3df,3pd) are far away from “chemical” accuracy (∼ 1 kcal/mol). In fact, p-ATV’s numerical value of PA=58.2 kcal/mol somewhat differs from ATV’s (and o-ATV’s) PA=61.5 kcal/mol, but if heterolytic O-H bond cleavage were to occur in p-ATV, it would rather occur at position 1-OH, which has a substantially smaller value PA=23.8 kcal/mol.

With regards to position 4-NH, the extra (5-)OH-group has a qualitatively different impact on the N-H bond cleavage of o-ATV and p-ATV. Notwithstanding the different values calculated (90.2 kcal/mol versus 89.3 kcal/mol), in the above vein we cannot soberly claim that H-atom abstraction from the NH-group is facilitated by the additional OH-group of o-ATV. However, the negative impact on the heterolytic N-H bond dissociation is significant. The o-ATV’s PA=49 kcal/mol is larger than the value PA=44.4 kcal/mol calculated for ATV. As of the heterolytic N-H bond dissociation, it is insensitively affected; the numerical difference between p-ATV’s PA=43.8 kcal/mol and ATV’s PA=44.4 kcal/mol obtained within B3LYP/6-31+G(d,p)/IEFPCM is too small to play a role in a sober analysis. Besides, similarly to what we said above, a heterolytic bond cleavage would occur at the lowest PA’s position 1-OH.

The really important effect brought about by the extra OH-group of the hydroxy metabolites is the homolytic bond dissociation at its position (5-OH). Our calculations demonstrate that this process is substantially less expensive energetically than H-atom donation from position 1-OH. The calculated BDE values for both o-ATV and p-ATV at this position are ∼ 77.5 kcal/mol versus the smallest value ∼ 91 kcal/mol for ATV at position 1-OH, respectively. Unlike the extremely similar homolytic bond dissociation, there is a certain difference between o-ATV’s and p-ATV’s heterolytic bond dissociation at position 5-OH, as expressed by the PA values (PA=34.4 kcal/mol ≠ PA=37.9 kcal/mol, respectively). However, it is unlikely that this difference in PA’s has practical consequences, again because the aforementioned values of PA are both comfortably larger than the lowest PA=23.8 kcal/mol at position 1-OH, a value that also characterizes the parent ATV molecule.

In Section 3.5 we will return to the practical implications of the above finding.

### 3.4. Alternative Approaches to the O-H and N-H Bond Strengths: Vibrational Frequencies and Bond Order Indices

The robustness of a single molecule diode fabricated using the scanning transmission microscopy (STM) break-junction technique [58,59] can be quantified by the maximum force that the junction subject to mechanical stretching can withstand. This rupture (pull-off) force *F* per molecule, which characterizes the strength of the chemical bond between electrodes and the terminal (anchoring) atom of the embedded molecule, can hardly be directly measured. Therefore, experimentalists use a simple mechanical model which relates *F* to the vibrational frequency of the pertaining stretching mode, which can be easily measured by infrared spectroscopy [60]. To exemplify, this is the Au – S stretching mode in benchmark nanojunctions wherein molecules are anchored via thiol groups on gold electrodes.

Applied to the present context, it is interesting to interrogate the relationship between BDE and the related stretching frequency. In the same vein, a stronger chemical X-Y bond is intuitively expected to have not only a larger BDE and a higher stretching frequency *ν*(X−Y) but also a shorter length and a larger bond order index.

Let us therefore examine the correlation of the aforementioned quantities in the presently considered molecules.

Infrared spectra calculated for ATV, o-OH-AVT, and p-ATV in methanol are depicted in Figure 7.

**Figure 7.**
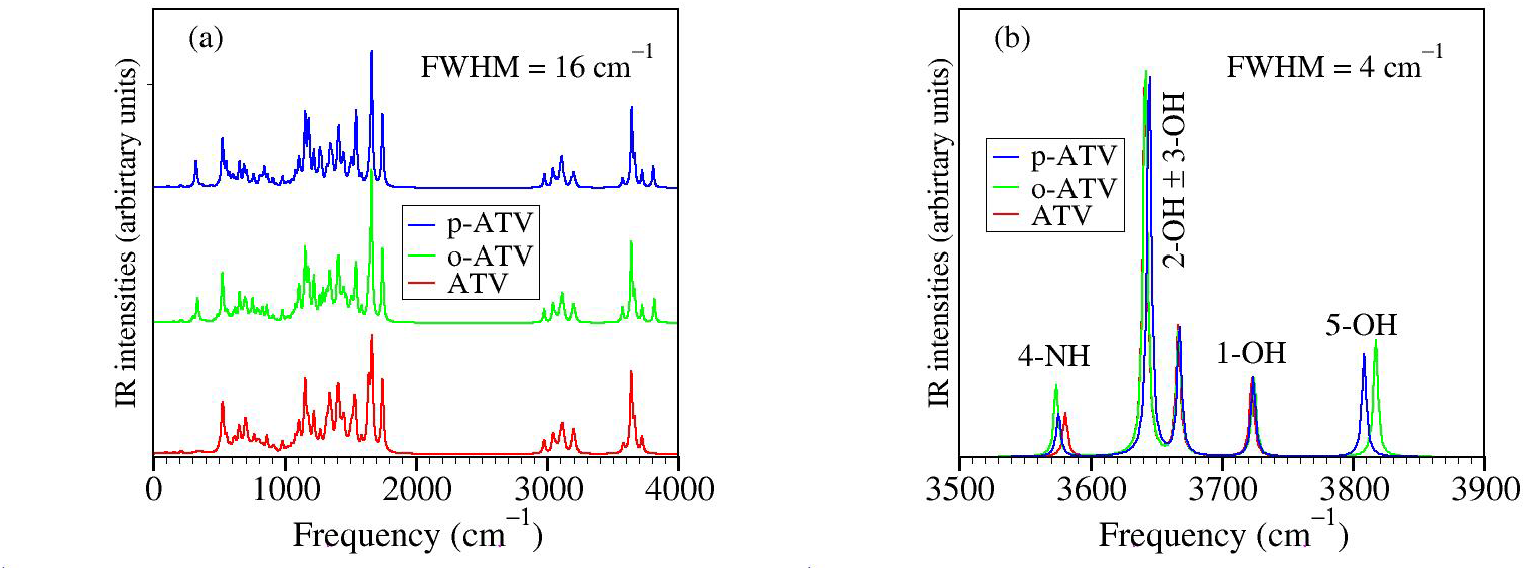
Infrared spectra calculated for ATV, o-OH-AVT, and p-ATV in methanol using Lorentzian convolution of full width at half maximum (FWHM) indicated in the inset: (a) in the whole range of frequency and (b) in the range where the O-H and N-H stretching modes are active. In all species, stretching modes of 2-OH and 3-OH groups appear as linear and antilinear vibrations rather than separated vibrational modes, and this may indicate that a more adequate optimization of the radicals generated by H-atom abstraction at these positions (which appear almost degenerate energetically, see pertaining BDE values in Table 4) should be done within a multi-reference framework.

**Table 4.**
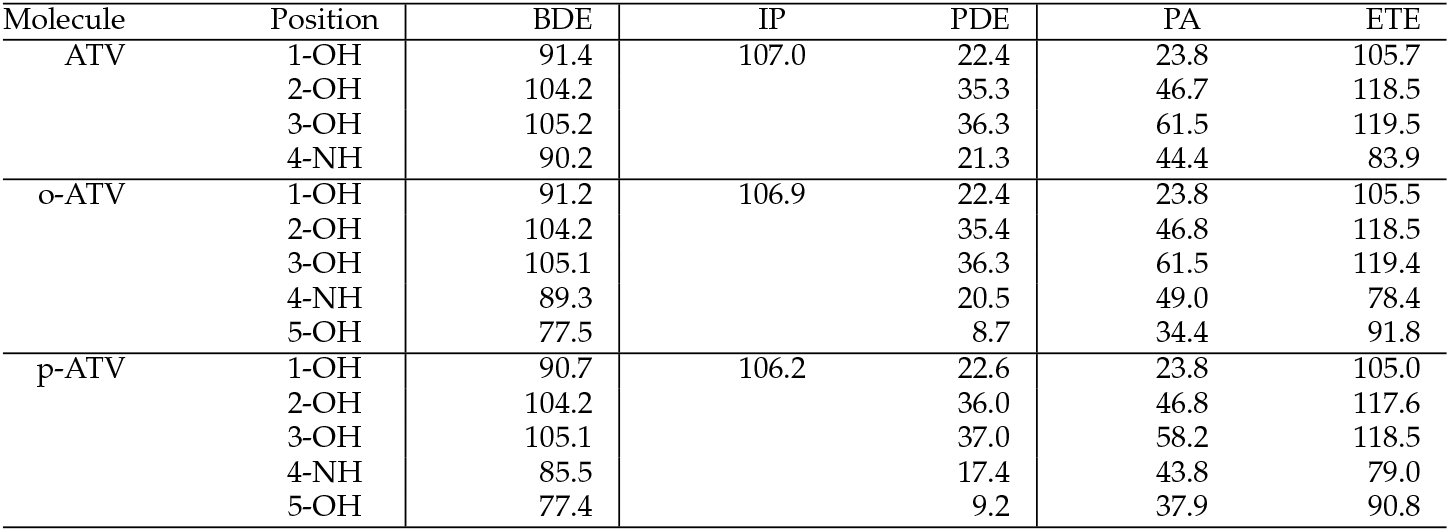
The enthalpies of reaction (in kcal/mol) needed to quantify the antioxidant activity of atorvastatin (ATV) and its ortho- and para-hydroxy metabolites (o-ATV, p-ATV).

**Table 5.**
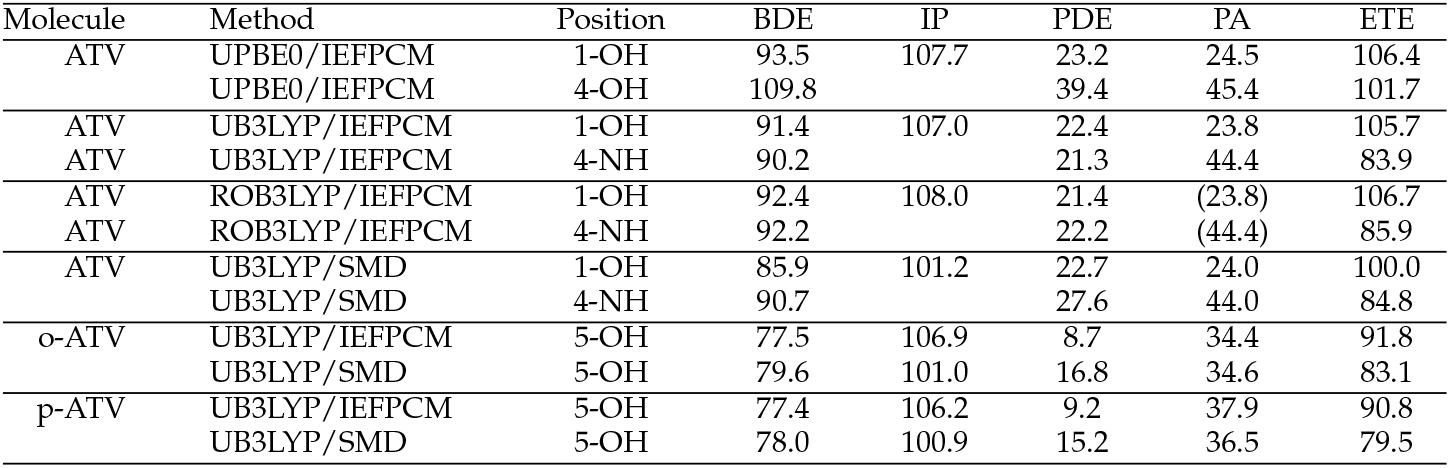
Enthalpies of reaction (in kcal/mol) computed for atorvastatin (ATV) using methods indicated above and 6-31+G(d,p) basis sets. There is no difference between unrestricted (UB3LYP) and restricted open shell (ROB3LYP) methods in calculating the PA values, and for this reason the pertaining value was written in parenthesis.

The behavior visible in Figure 7b is surprising for several reasons, e.g.:

i. although the BDE of ATV and its metabolites at position 1-OH is lower than at positions 2-OH and 3-OH, the streching mode at position 1-OH has a higher frequency than at positions 2-OH and 3-OH;
ii. although o-ATV and p-ATV have at position 5-OH a smaller BDE than for all OH-positions of the parent ATV, the 5-OH stretching mode of the metabolites is higher than those of all O-H streching mode of ATV;
iii. although o-ATV’s and ATV’s N-H BDE are equal, the frequency of the N-H of the former is smaller than that of the latter;
iv. although o-ATV’s BDE and p-ATV’s BDE are different, their N-H streching modes have the same frequency;
v. although o-ATV and p-ATV have equal BDE at position 5-OH, the o-ATV’s O-H streching frequency is higher than that of p-ATV.

Counter-intuitive aspects of the relationship BDE versus *ν* are visualized in Figures 8a and 9a.

**Figure 8.**
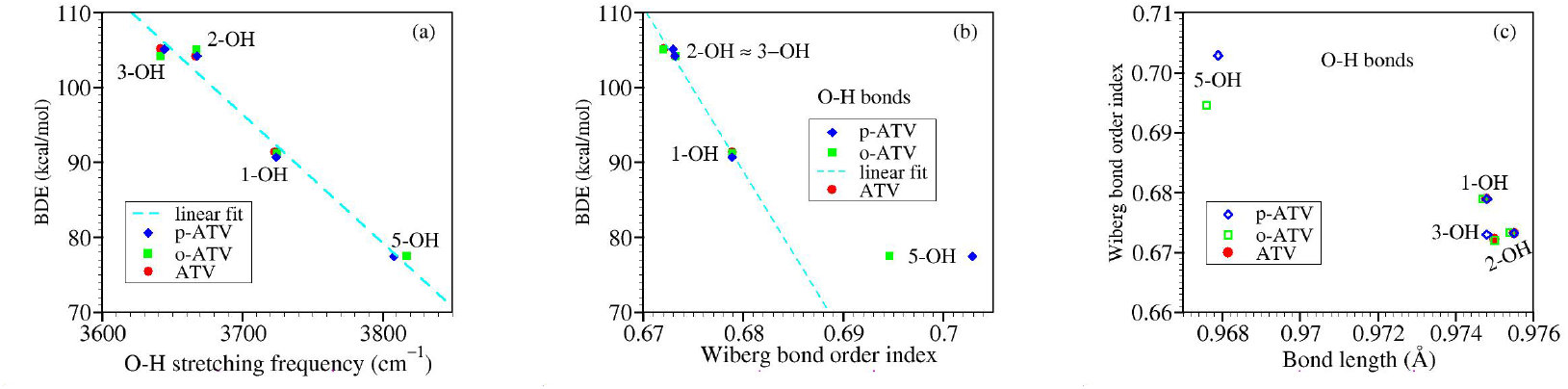
Results for OH groups of atorvastatin (ATV) and its metabolites o-ATV and p-ATV: (a) bond dissociation energies versus O-H stretching frequencies; (b) bond dissociation energies versus Wiberg bond order indices; (c) Wiberg bond order indices versus bond lengths.

Let us now switch to bond order indices. Our results are collected in Table 6, and Figures 8 and 9.

**Table 6.** Wiberg bond order indices, bond lengths (in Å), vibrational frequencies (in cm−1, and bond dissociation energies BDE (in kcal/mol) for atorvastatin and its metabolites.

| Molecule | Position | Wiberg | Length | BDE | $\nu$ |
| --- | --- | --- | --- | --- | --- |
| ATV | 1-OH | 0.6789 | 0.9748 | 91.4 | 3723.2 |
|  | 2-OH | 0.6732 | 0.9755 | 104.2 | 3667.0 |
|  | 3-OH | 0.6721 | 0.9750 | 105.2 | 3641.6 |
|  | 4-NH | 0.7562 | 1.0147 | 90.2 | 3581.0 |
| o-ATV | 1-OH | 0.6789 | 0.9747 | 91.2 | 3724.8 |
|  | 2-OH | 0.6733 | 0.9754 | 104.2 | 3641.7 |
|  | 3-OH | 0.6720 | 0.9750 | 105.1 | 3667.5 |
|  | 4-NH | 0.7454 | 1.0151 | 89.3 | 3574.2 |
|  | 5-OH | 0.6946 | 0.9676 | 77.5 | 3817.1 |
| p-ATV | 1-OH | 0.6789 | 0.9748 | 90.7 | 3724.2 |
|  | 2-OH | 0.6732 | 0.9755 | 104.2 | 3667.9 |
|  | 3-OH | 0.6730 | 0.9748 | 105.1 | 3644.7 |
|  | 4-NH | 0.7550 | 1.0149 | 85.5 | 3576.0 |
|  | 5-OH | 0.7029 | 0.9679 | 77.4 | 3808.4 |

**Figure 9.**
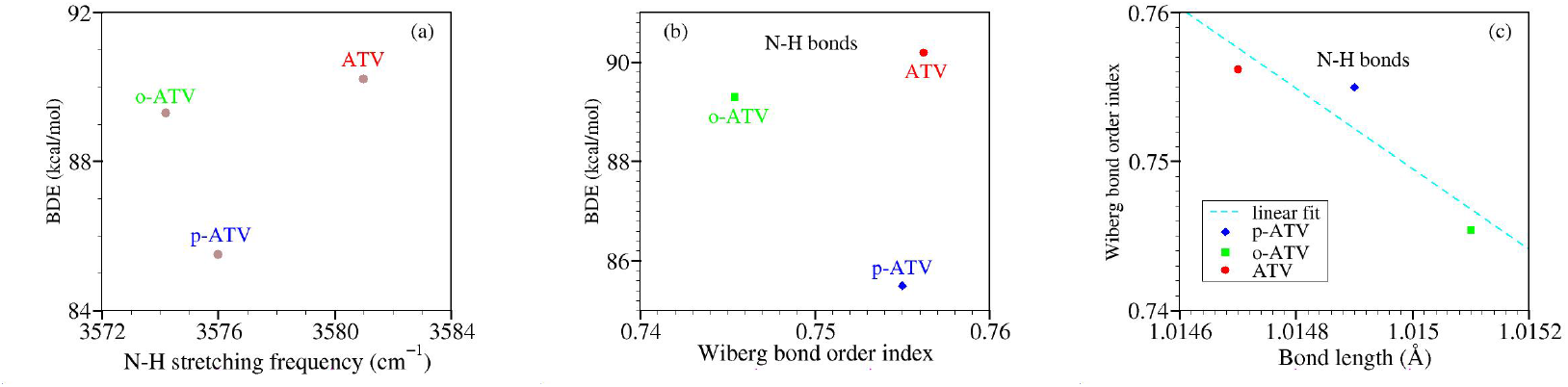
Results similar to Figure 8 but for NH groups.

**Figure 10.**
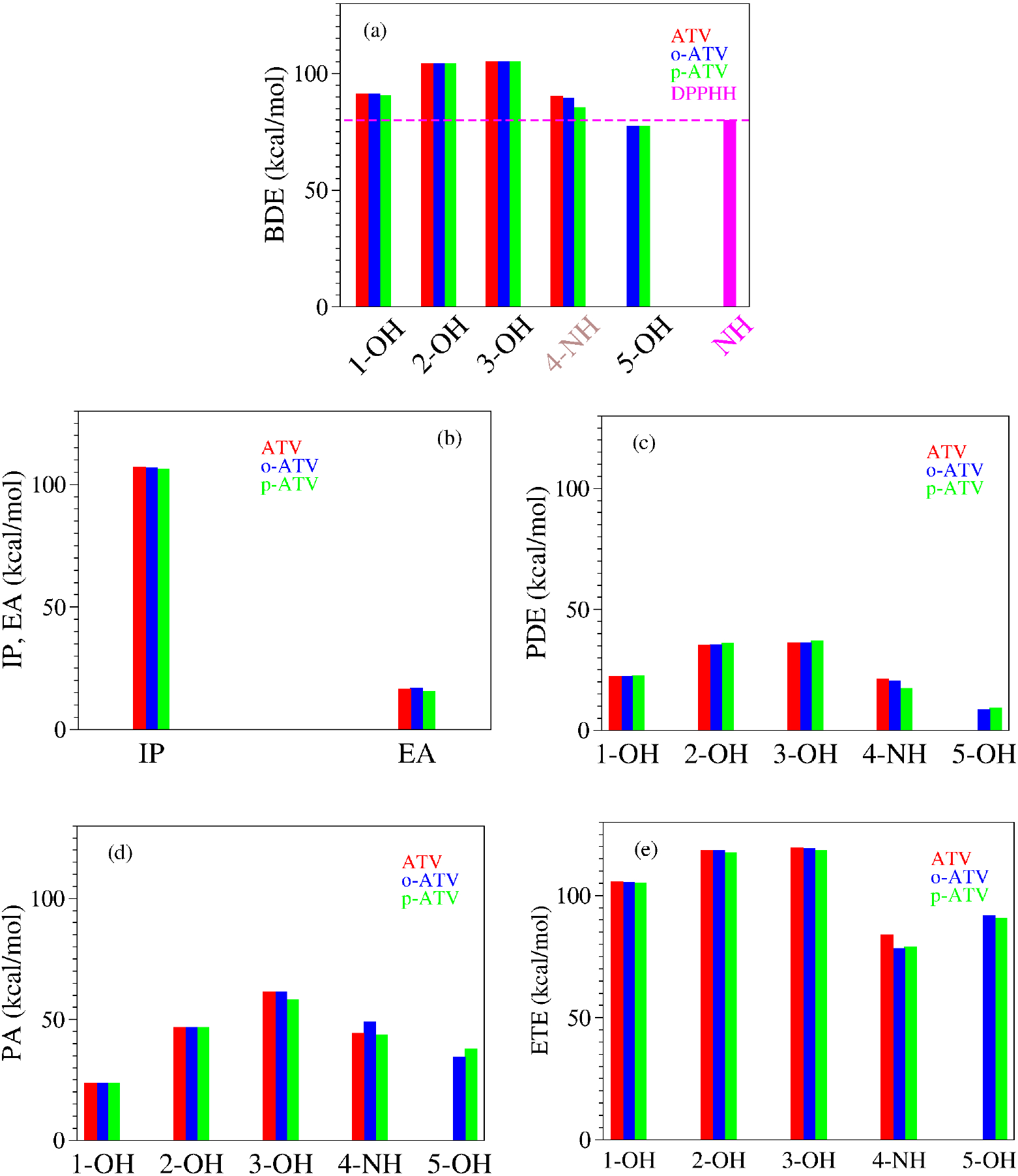
Enthalpies of reaction quantifying the antioxidant activity of atorvastatin (ATV) and its ortho-(o-ATV) and para- (p-ATV) hydroxy metabolites: (a) bond dissociation; (b) ionization and electron attachment; (c) proton detachment; (d) proton affinity; (e) electron transfer. The additional information for the DPPH• radical in panel (a) depicts why o-ATV and p-ATV can scavenge this radical while the parent ATV cannot.

To reiterate, based on straightforward chemical intuition, it would be obvious to expect that stronger chemical bonds (larger BDE’s) possess larger bond order indices. Figure 8b depicts that for the O-H bonds of ATV, o-ATV, and p-ATV just the opposite holds true: larger BDE’s justly correspond to smaller bond order indices. As for their N-H bonds, Figure 8b reveals that the dependence is even nonmonotonic.

To avoid misunderstanding, a clarification is in order before ending this analysis. What chemical intuition in the above example should not overlook is that a pair of atoms X and Y forming an X-Y chemical bond, do not merely interact with each other but also with the neighboring atoms in the molecular surrounding. This is also why a simple (exponential [61]) relationship between bond order indices and bond lengths can hold, e.g., for homologous molecular series [62], but cannot not hold in general; otherwise one arrives at comparing apples with oranges. Figures 8 and 9 illustrate this again using the values of Table 6. BDE values corresponding to different O-H bonds of a given molecule differ from each other depending on the specific chemical environment. These differences can be visualized by inspecting the spin density landscape of the various radicals (Figures 1, 3, 4, and 5). The stronger the delocalization in a radical, the easier is its formation, and the lower is the corresponding BDE value. Inspection of Figures 1b and c makes it clear, e.g., why ATV’s BDE at position 3-OH is higher than that at position 1-OH.

### 3.5. Assessing the Radical Scavenging Activity. A Specific Example

Discussion on free radical scavenging and dominant antioxidant mechanism is very often couched by comparing among themselves values the enthalpies characterizing the HAT, SET-PL, and SPLET of the specific antioxidant(s) under investigation. Every now and then publications conclude, e.g., that SPLET is the dominant pathway because a certain antioxidant has a “small” PA value or a PA substantially smaller than BDE, or that SET-PL prevails because of the small IP value. However, it is worth emphasizing that, along with the antioxidant’s properties, a proper evaluation of the antioxidant activity should mandatory consider the specific properties of the radicals to be eliminated (neutralized).

The small value BDE≈ 77.5 kcal/mol for o-ATV and p-ATV, substantially smaller than the smallest value (BDE=90.2 kcal/mol) of the parent ATV, is perhaps the most appealing result reported in Section 3.3. Still, the “small” value mentioned above does not demonstrate *per se* the fact anticipated in Introduction, namely that o-ATV and p-ATV can scavenge can scavenge the ubiquitously employed 1,1-diphenyl-2-picrylhydrazyl (DPPH•) radical, while the parent ATV cannot.

To demonstrate this, one should mandatory consider the pertaining DPPH• property, namely the enthalpy release in DPPH•’s neutralization (H-atom affinity)

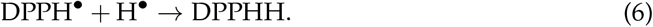

Because it amounts to 80 kcal/mol [63], e.g., the reaction

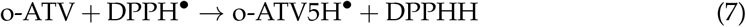

is exothermic. H-atom abstraction from position 5-OH of o-ATV (or p-ATV) costs ∼ 77.5 kcal/mol, a value lower that the enthalpy release of 80 kcal/mol [63] in the neutralization of the DPPH• radical, and this makes the HAT mechanism thermodynamically allowed. Rephrasing, because the BDE of the N – H bond of DPPHH is 80 kcal/mol [63], o-ATV (and p-ATV) can scavenge the DPPH• radical through donating the H-atom at position 5-OH. On the contrary, the parent ATV cannot. The lowest ATV’s BDE (at position 1-OH) amounts to 90.2 kcal/mol (Table 4), so the HAT pathway is forbidden.

To conclude, we have presented above the first theoretical explanation of the experimental fact [8] that the antioxidant properties of atorvastatin ortho- and para-hydroxy metabolites differ from those of ATV.

By and large, there is a consensus in the literature that HAT is a possible (or even preferred) antioxidant mechanism in the gases phase but not in polar protic solvents like the presently considered methanol. In this vein, the natural question that arises is: can o-ATV and p-ATV scavenge the DPPH• radical in methanol also via SPLET? Can HAT and SPLET coexist? While the large IP (Table 4) give little chances to an SET-PT pathway, SPLET would a priori be conceivable in view of the “small” value of PA, which is, although not smaller than that of ascorbic acid (as incorrectly [6] claimed [5]) at least not much larger than the latter (23.8 kcal/mol for ATV’s versus 20.5 kcal/mol for ascorbic acid, see ref. [6]).

In fact, Table 4 implicitly gives the *negative* answer to this question. If o-ATV and p-ATV could scavenge DPPH• via SPLET, then (contrary to experiment [8]) the parent ATV could also do the job; the most favored deprotonation, implying the same enthalpy PA=23.8 kcal/mol, occurs both for ATV and its metabolites at the same 1-OH position, where furthermore the similar spin density landscapes (compare Figure 1b with Figure 3) indicate a similar chemical reactivity.

Still, let us remain in the realm of theory and demonstrate why neither o-ATV nor p-ATV or ATV can scavenge DPPH• in methanol. To this aim suffice it to consider the first step of SPLET

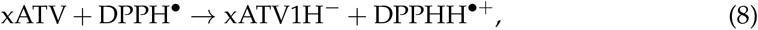

where x means “o-”, “p-”, or “nothing”. Straightforward manipulation allows to express the enthalpy of this reaction as follows

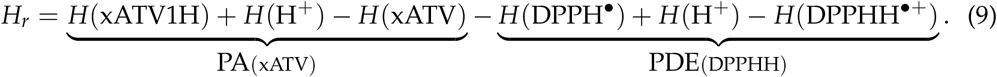

Notice that the second brace in equation (9) corresponds to the proton abstraction from the cation DPPHH^•+^ of the neutralized free radical DPPHH, or alternatively, the PDE pertaining to the neutralized free radical DPPHH (cf. equation (3b)).

Equation (9) reveals that, to be thermodynamically allowed, the first SPLET step requires

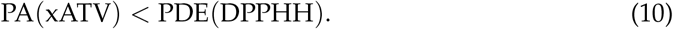

Our calculations yielded PDE(DPPHH) = 3.9 kcal/mol, a value that is not larger (as the case if the first SPLET step was allowed) but smaller than PA(xATV) = 23.8 kcal/mol. It now becomes clear why neither ATV, nor o-ATV or p-ATV can scavenge the DPPH• radical via SPLET. Their “small” PA is not small enough to fulfill equation (10).

### 4. Conclusion

We believe that the present demonstration that atorvastatin ortho- and para-hydroxy metabolites can scavenge the DPPH• through donating the H-atom at the position of their extra group (5-OH), which is impossible in the parent ATV, is important not only because it theoretically explains for the first time a behavior revealed in experiment [8] but also because, from a general perspective, it provides further insight into the structure–activity relationship (SAR).

By working out a specific example (Section 3.5) — an analysis that can be straight-forwardly extended to other cases—, we drew attention that an adequate approach to antioxidant’s potency should mandatory account for the thermodynamic properties of the free radicals. Equation (10) expresses a general necessary condition for thermodynamicallyy allowed SPLET, and its application to specific cases may reveal that, even in polar solvents, free radical scavenging via this is pathway forbidden not only for ATV-based species.

In addition, our study emphasize that, while important, e.g., for modeling the temporal evolution of various molecular species interacting among themselves in a given chemical environment [62,64], the global chemical reactivity indices have no direct relevance for antioxidation. Recall that we saw in Section 3.2 that quantitative differences of ATV’s o-ATV’s, and p-ATV’s global chemical reactivity indices are minor. Furthermore, if qualitative differences in these indices were important, then, contrary to Sections 3.3 and 3.5, o-ATV would have antioxidant properties similar to ATV rather than to p-ATV.

Last but not least, from the perspective of fundamental science, we found (Section 3.4) that properties like bond dissociation enthalpy, bond order index, bond length, and bond stretching frequency, expected after all to represent alternatives in quantifying the bond strength, are by no means correlated according to naive intuition. This finding calls for further quantum chemical efforts aiming at comprehensively characterizing ATV’s, that inherently remained beyond the scope of this study focused on ATV’s antioxidant activity. Finally, the presently reported counter-intutitve relationship between bond stretching frequency and bond strength should also be a word of caution for other communities; for example, for the molecular electronics community, wherein bond stretching frequencies (conveniently obtained via infrared spectroscopy) are used to estimate (pull-off) forces that cause the rupture of a junction subject to mechanical stretching [65].

## Acknowledgments

The author is much indebted to Ederley Vélez Ortiz for providing valuable details related to her recent work [5]. Financial support from the German Research Foundation (DFG Grant No. BA 1799/3-2) in the initial stage of this work and computational support by the state of Baden-Württemberg through bwHPC and the German Research Foundation through Grant No. INST 40/575-1 FUGG (bwUniCluster 2.0, bwForCluster/MLS&WISO 2.0, and JUSTUS 2.0 cluster) are gratefully acknowledged.

## Appendix A

**Table A1.**
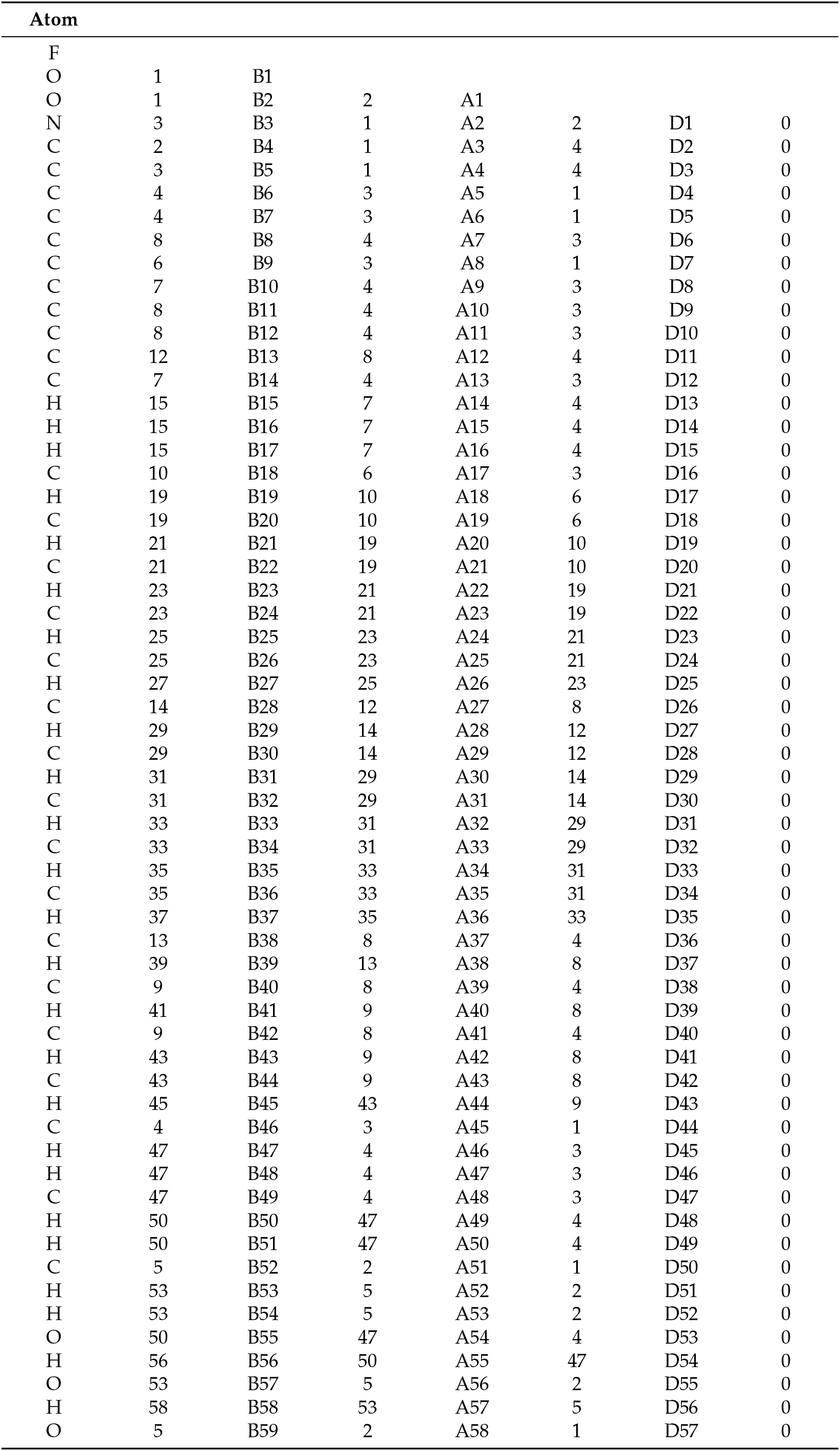
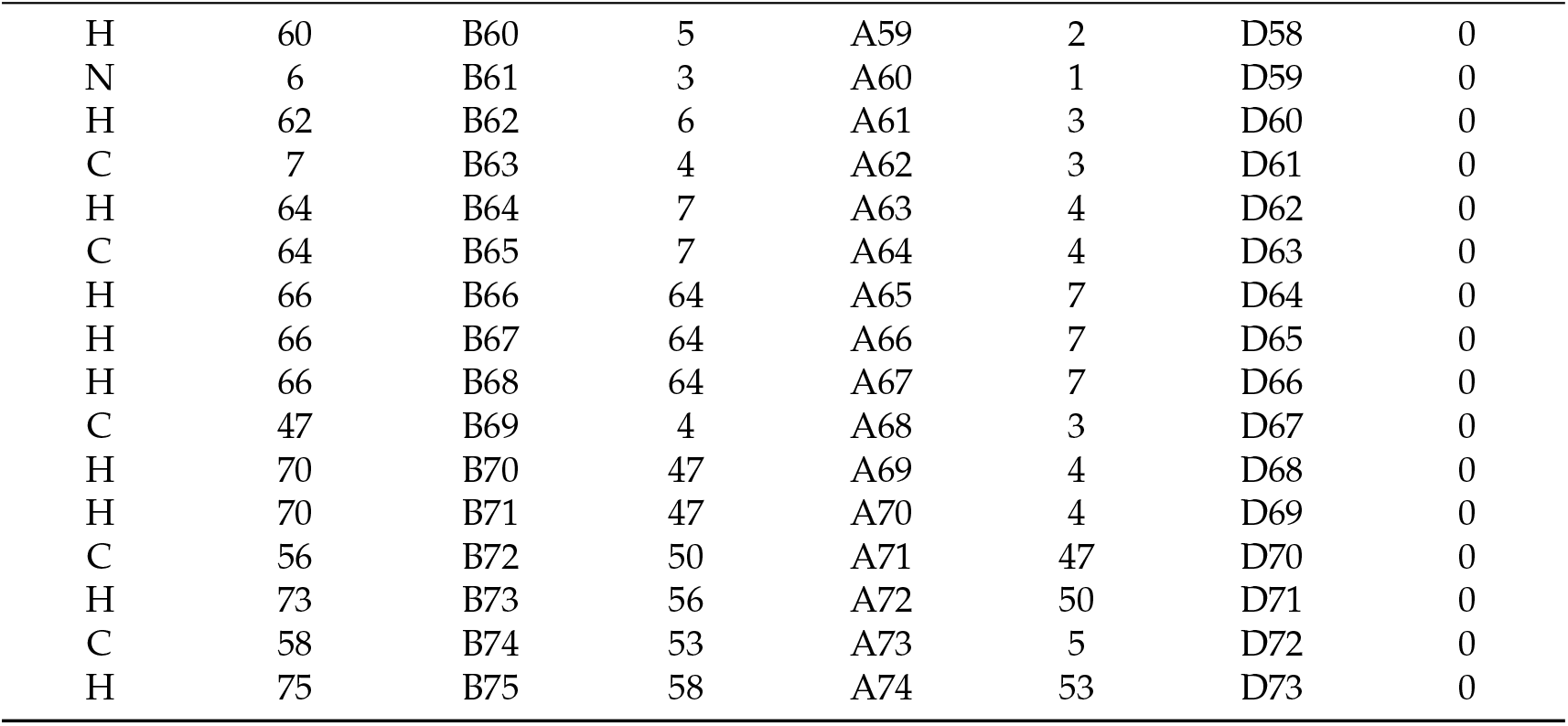
Z−matrix of ATV.

| Atom |  |  |  |  |  |  |  |
| --- | --- | --- | --- | --- | --- | --- | --- |
| F |  |  |  |  |  |  |  |
| O | 1 | B1 |  |  |  |  |  |
| O | 1 | B2 | 2 | A1 |  |  |  |
| N | 3 | B3 | 1 | A2 | 2 | D1 | 0 |
| C | 2 | B4 | 1 | A3 | 4 | D2 | 0 |
| C | 3 | B5 | 1 | A4 | 4 | D3 | 0 |
| C | 4 | B6 | 3 | A5 | 1 | D4 | 0 |
| C | 4 | B7 | 3 | A6 | 1 | D5 | 0 |
| C | 8 | B8 | 4 | A7 | 3 | D6 | 0 |
| C | 6 | B9 | 3 | A8 | 1 | D7 | 0 |
| C | 7 | B10 | 4 | A9 | 3 | D8 | 0 |
| C | 8 | B11 | 4 | A10 | 3 | D9 | 0 |
| C | 8 | B12 | 4 | A11 | 3 | D10 | 0 |
| C | 12 | B13 | 8 | A12 | 4 | D11 | 0 |
| C | 7 | B14 | 4 | A13 | 3 | D12 | 0 |
| H | 15 | B15 | 7 | A14 | 4 | D13 | 0 |
| H | 15 | B16 | 7 | A15 | 4 | D14 | 0 |
| H | 15 | B17 | 7 | A16 | 4 | D15 | 0 |
| C | 10 | B18 | 6 | A17 | 3 | D16 | 0 |
| H | 19 | B19 | 10 | A18 | 6 | D17 | 0 |
| C | 19 | B20 | 10 | A19 | 6 | D18 | 0 |
| H | 21 | B21 | 19 | A20 | 10 | D19 | 0 |
| C | 21 | B22 | 19 | A21 | 10 | D20 | 0 |
| H | 23 | B23 | 21 | A22 | 19 | D21 | 0 |
| C | 23 | B24 | 21 | A23 | 19 | D22 | 0 |
| H | 25 | B25 | 23 | A24 | 21 | D23 | 0 |
| C | 25 | B26 | 23 | A25 | 21 | D24 | 0 |
| H | 27 | B27 | 25 | A26 | 23 | D25 | 0 |
| C | 14 | B28 | 12 | A27 | 8 | D26 | 0 |
| H | 29 | B29 | 14 | A28 | 12 | D27 | 0 |
| C | 29 | B30 | 14 | A29 | 12 | D28 | 0 |
| H | 31 | B31 | 29 | A30 | 14 | D29 | 0 |
| C | 31 | B32 | 29 | A31 | 14 | D30 | 0 |
| H | 33 | B33 | 31 | A32 | 29 | D31 | 0 |
| C | 33 | B34 | 31 | A33 | 29 | D32 | 0 |
| H | 35 | B35 | 33 | A34 | 31 | D33 | 0 |
| C | 35 | B36 | 33 | A35 | 31 | D34 | 0 |
| H | 37 | B37 | 35 | A36 | 33 | D35 | 0 |
| C | 13 | B38 | 8 | A37 | 4 | D36 | 0 |
| H | 39 | B39 | 13 | A38 | 8 | D37 | 0 |
| C | 9 | B40 | 8 | A39 | 4 | D38 | 0 |
| H | 41 | B41 | 9 | A40 | 8 | D39 | 0 |
| C | 9 | B42 | 8 | A41 | 4 | D40 | 0 |
| H | 43 | B43 | 9 | A42 | 8 | D41 | 0 |
| C | 43 | B44 | 9 | A43 | 8 | D42 | 0 |
| H | 45 | B45 | 43 | A44 | 9 | D43 | 0 |
| C | 4 | B46 | 3 | A45 | 1 | D44 | 0 |
| H | 47 | B47 | 4 | A46 | 3 | D45 | 0 |
| H | 47 | B48 | 4 | A47 | 3 | D46 | 0 |
| C | 47 | B49 | 4 | A48 | 3 | D47 | 0 |
| H | 50 | B50 | 47 | A49 | 4 | D48 | 0 |
| H | 50 | B51 | 47 | A50 | 4 | D49 | 0 |
| C | 5 | B52 | 2 | A51 | 1 | D50 | 0 |
| H | 53 | B53 | 5 | A52 | 2 | D51 | 0 |
| H | 53 | B54 | 5 | A53 | 2 | D52 | 0 |
| O | 50 | B55 | 47 | A54 | 4 | D53 | 0 |
| H | 56 | B56 | 50 | A55 | 47 | D54 | 0 |
| O | 53 | B57 | 5 | A56 | 2 | D55 | 0 |
| H | 58 | B58 | 53 | A57 | 5 | D56 | 0 |
| O | 5 | B59 | 2 | A58 | 1 | D57 | 0 |
| H | 60 | B60 | 5 | A59 | 2 | D58 | 0 |
| N | 6 | B61 | 3 | A60 | 1 | D59 | 0 |
| H | 62 | B62 | 6 | A61 | 3 | D60 | 0 |
| C | 7 | B63 | 4 | A62 | 3 | D61 | 0 |
| H | 64 | B64 | 7 | A63 | 4 | D62 | 0 |
| C | 64 | B65 | 7 | A64 | 4 | D63 | 0 |
| H | 66 | B66 | 64 | A65 | 7 | D64 | 0 |
| H | 66 | B67 | 64 | A66 | 7 | D65 | 0 |
| H | 66 | B68 | 64 | A67 | 7 | D66 | 0 |
| C | 47 | B69 | 4 | A68 | 3 | D67 | 0 |
| H | 70 | B70 | 47 | A69 | 4 | D68 | 0 |
| H | 70 | B71 | 47 | A70 | 4 | D69 | 0 |
| C | 56 | B72 | 50 | A71 | 47 | D70 | 0 |
| H | 73 | B73 | 56 | A72 | 50 | D71 | 0 |
| C | 58 | B74 | 53 | A73 | 5 | D72 | 0 |
| H | 75 | B75 | 58 | A74 | 53 | D73 | 0 |

**Table A2.**
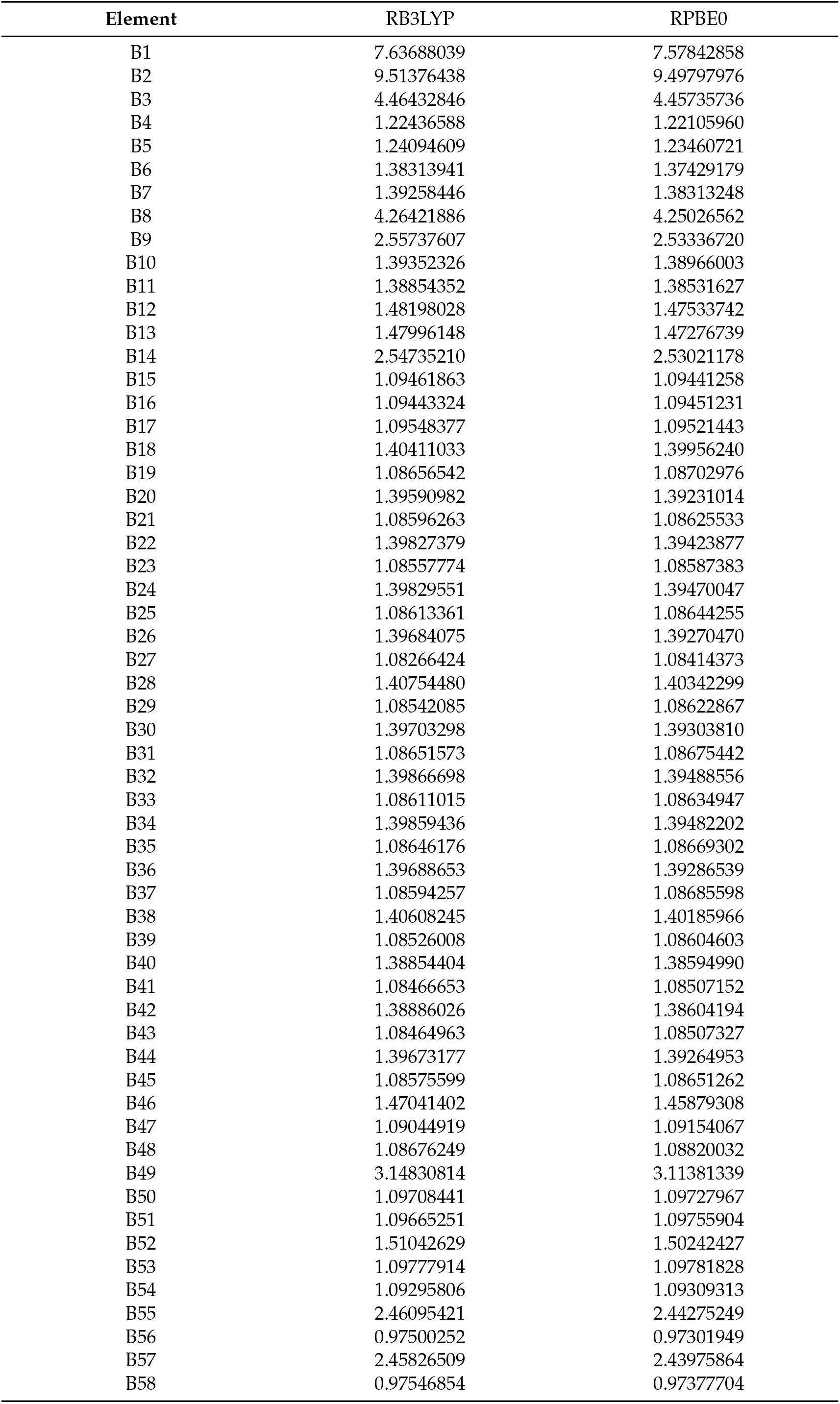
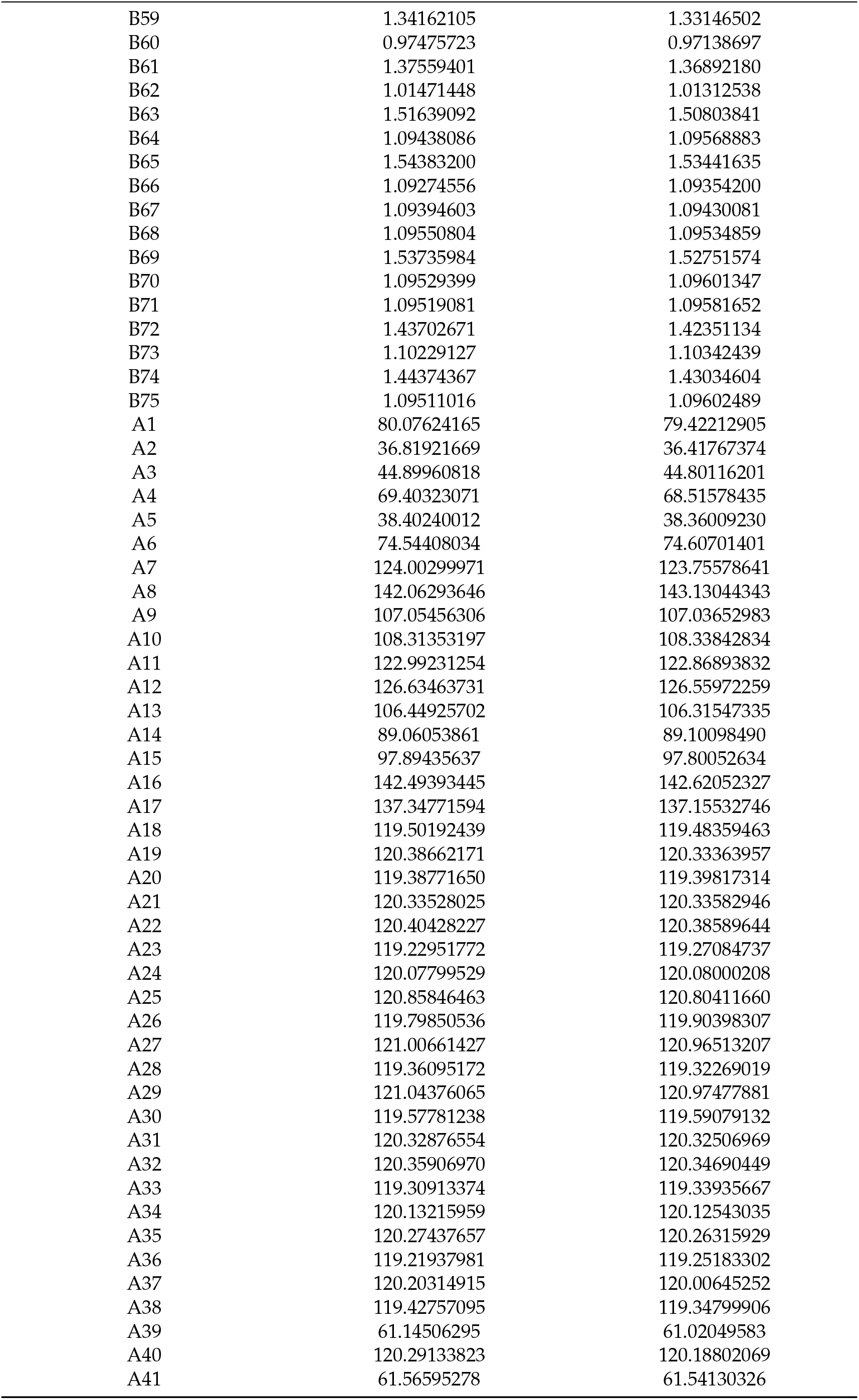
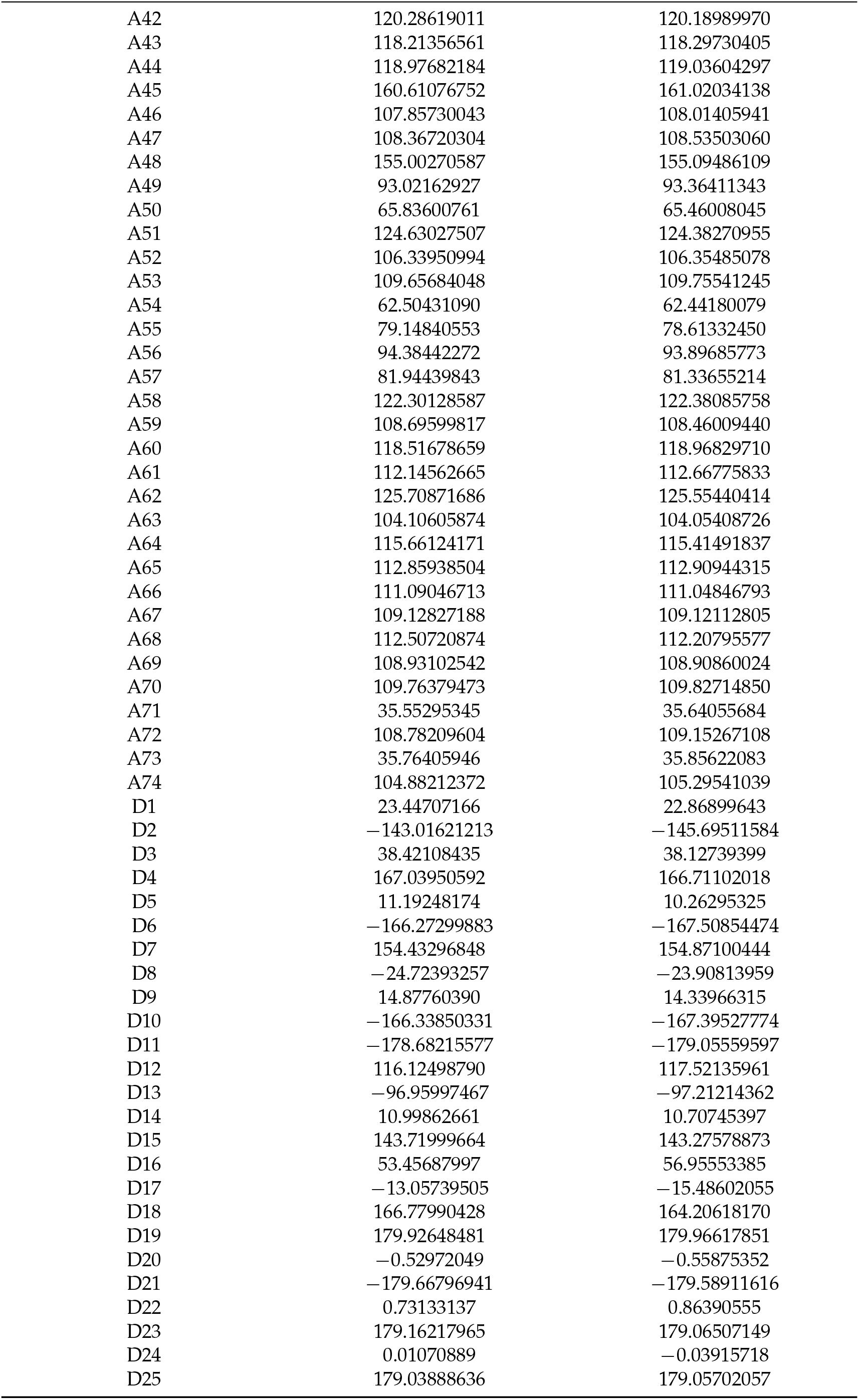
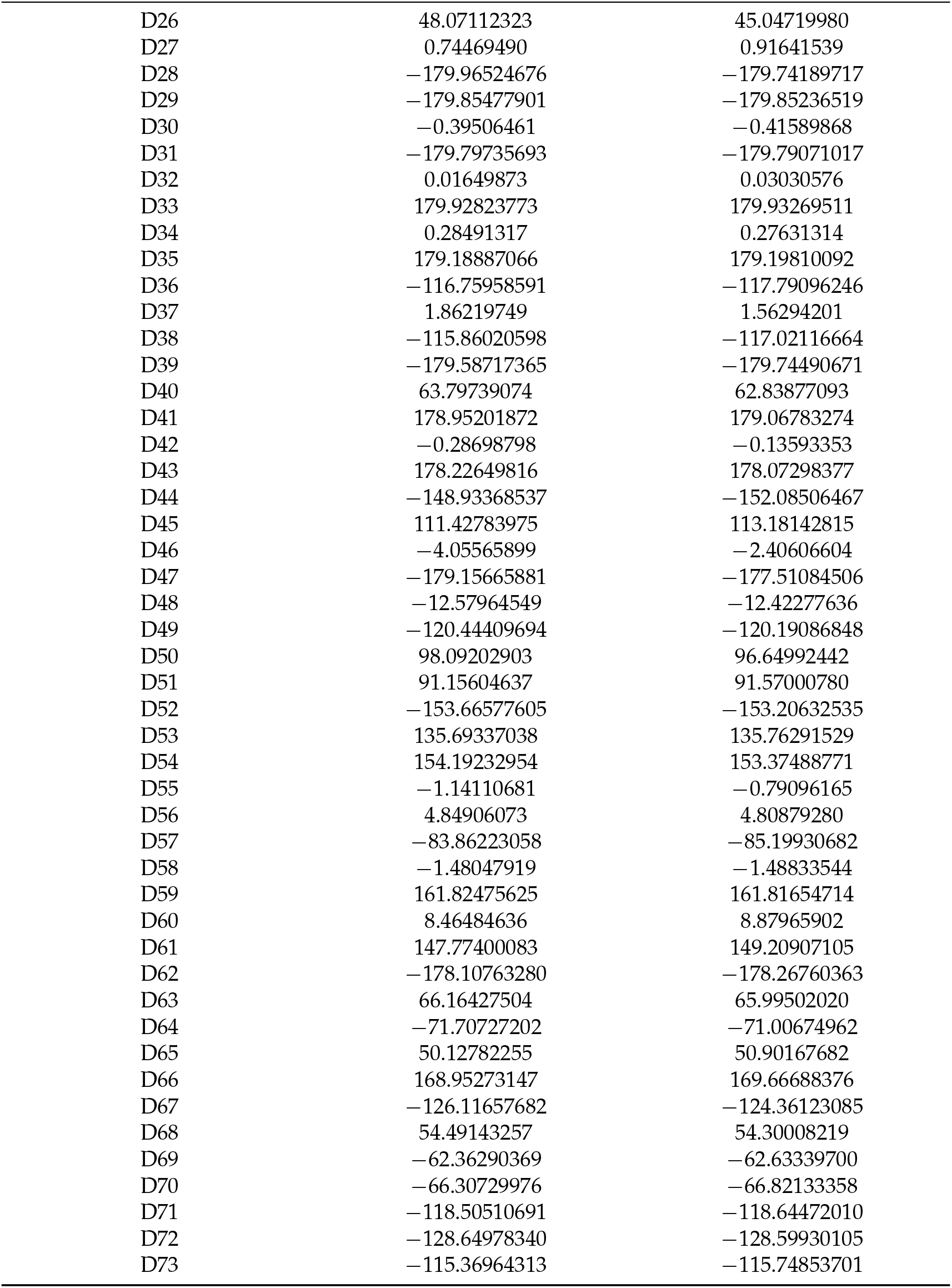
Elements of the Z−matrix of ATV in methanol optimized as indicated below using 6-31+G/d,p) basis sets.

| Element | RB3LYP | RPBE0 |
| --- | --- | --- |
| B1 | 7.63688039 | 7.57842858 |
| B2 | 9.51376438 | 9.49797976 |
| B3 | 4.46432846 | 4.45735736 |
| B4 | 1.22436588 | 1.22105960 |
| B5 | 1.24094609 | 1.23460721 |
| B6 | 1.38313941 | 1.37429179 |
| B7 | 1.39258446 | 1.38313248 |
| B8 | 4.26421886 | 4.25026562 |
| B9 | 2.55737607 | 2.53336720 |
| B10 | 1.39352326 | 1.38966003 |
| B11 | 1.38854352 | 1.38531627 |
| B12 | 1.48198028 | 1.47533742 |
| B13 | 1.47996148 | 1.47276739 |
| B14 | 2.54735210 | 2.53021178 |
| B15 | 1.09461863 | 1.09441258 |
| B16 | 1.09443324 | 1.09451231 |
| B17 | 1.09548377 | 1.09521443 |
| B18 | 1.40411033 | 1.39956240 |
| B19 | 1.08656542 | 1.08702976 |
| B20 | 1.39590982 | 1.39231014 |
| B21 | 1.08596263 | 1.08625533 |
| B22 | 1.39827379 | 1.39423877 |
| B23 | 1.08557774 | 1.08587383 |
| B24 | 1.39829551 | 1.39470047 |
| B25 | 1.08613361 | 1.08644255 |
| B26 | 1.39684075 | 1.39270470 |
| B27 | 1.08266424 | 1.08414373 |
| B28 | 1.40754480 | 1.40342299 |
| B29 | 1.08542085 | 1.08622867 |
| B30 | 1.39703298 | 1.39303810 |
| B31 | 1.08651573 | 1.08675442 |
| B32 | 1.39866698 | 1.39488556 |
| B33 | 1.08611015 | 1.08634947 |
| B34 | 1.39859436 | 1.39482202 |
| B35 | 1.08646176 | 1.08669302 |
| B36 | 1.39688653 | 1.39286539 |
| B37 | 1.08594257 | 1.08685598 |
| B38 | 1.40608245 | 1.40185966 |
| B39 | 1.08526008 | 1.08604603 |
| B40 | 1.38854404 | 1.38594990 |
| B41 | 1.08466653 | 1.08507152 |
| B42 | 1.38886026 | 1.38604194 |
| B43 | 1.08464963 | 1.08507327 |
| B44 | 1.39673177 | 1.39264953 |
| B45 | 1.08575599 | 1.08651262 |
| B46 | 1.47041402 | 1.45879308 |
| B47 | 1.09044919 | 1.09154067 |
| B48 | 1.08676249 | 1.08820032 |
| B49 | 3.14830814 | 3.11381339 |
| B50 | 1.09708441 | 1.09727967 |
| B51 | 1.09665251 | 1.09755904 |
| B52 | 1.51042629 | 1.50242427 |
| B53 | 1.09777914 | 1.09781828 |
| B54 | 1.09295806 | 1.09309313 |
| B55 | 2.46095421 | 2.44275249 |
| B56 | 0.97500252 | 0.97301949 |
| B57 | 2.45826509 | 2.43975864 |
| B58 | 0.97546854 | 0.97377704 |
| B59 | 1.34162105 | 1.33146502 |
| B60 | 0.97475723 | 0.97138697 |
| B61 | 1.37559401 | 1.36892180 |
| B62 | 1.01471448 | 1.01312538 |
| B63 | 1.51639092 | 1.50803841 |
| B64 | 1.09438086 | 1.09568883 |
| B65 | 1.54383200 | 1.53441635 |
| B66 | 1.09274556 | 1.09354200 |
| B67 | 1.09394603 | 1.09430081 |
| B68 | 1.09550804 | 1.09534859 |
| B69 | 1.53735984 | 1.52751574 |
| B70 | 1.09529399 | 1.09601347 |
| B71 | 1.09519081 | 1.09581652 |
| B72 | 1.43702671 | 1.42351134 |
| B73 | 1.10229127 | 1.10342439 |
| B74 | 1.44374367 | 1.43034604 |
| B75 | 1.09511016 | 1.09602489 |
| A1 | 80.07624165 | 79.42212905 |
| A2 | 36.81921669 | 36.41767374 |
| A3 | 44.89960818 | 44.80116201 |
| A4 | 69.40323071 | 68.51578435 |
| A5 | 38.40240012 | 38.36009230 |
| A6 | 74.54408034 | 74.60701401 |
| A7 | 124.00299971 | 123.75578641 |
| A8 | 142.06293646 | 143.13044343 |
| A9 | 107.05456306 | 107.03652983 |
| A10 | 108.31353197 | 108.33842834 |
| A11 | 122.99231254 | 122.86893832 |
| A12 | 126.63463731 | 126.55972259 |
| A13 | 106.44925702 | 106.31547335 |
| A14 | 89.06053861 | 89.10098490 |
| A15 | 97.89435637 | 97.80052634 |
| A16 | 142.49393445 | 142.62052327 |
| A17 | 137.34771594 | 137.15532746 |
| A18 | 119.50192439 | 119.48359463 |
| A19 | 120.38662171 | 120.33363957 |
| A20 | 119.38771650 | 119.39817314 |
| A21 | 120.33528025 | 120.33582946 |
| A22 | 120.40428227 | 120.38589644 |
| A23 | 119.22951772 | 119.27084737 |
| A24 | 120.07799529 | 120.08000208 |
| A25 | 120.85846463 | 120.80411660 |
| A26 | 119.79850536 | 119.90398307 |
| A27 | 121.00661427 | 120.96513207 |
| A28 | 119.36095172 | 119.32269019 |
| A29 | 121.04376065 | 120.97477881 |
| A30 | 119.57781238 | 119.59079132 |
| A31 | 120.32876554 | 120.32506969 |
| A32 | 120.35906970 | 120.34690449 |
| A33 | 119.30913374 | 119.33935667 |
| A34 | 120.13215959 | 120.12543035 |
| A35 | 120.27437657 | 120.26315929 |
| A36 | 119.21937981 | 119.25183302 |
| A37 | 120.20314915 | 120.00645252 |
| A38 | 119.42757095 | 119.34799906 |
| A39 | 61.14506295 | 61.02049583 |
| A40 | 120.29133823 | 120.18802069 |
| A41 | 61.56595278 | 61.54130326 |
| A42 | 120.28619011 | 120.18989970 |
| A43 | 118.21356561 | 118.29730405 |
| A44 | 118.97682184 | 119.03604297 |
| A45 | 160.61076752 | 161.02034138 |
| A46 | 107.85730043 | 108.01405941 |
| A47 | 108.36720304 | 108.53503060 |
| A48 | 155.00270587 | 155.09486109 |
| A49 | 93.02162927 | 93.36411343 |
| A50 | 65.83600761 | 65.46008045 |
| A51 | 124.63027507 | 124.38270955 |
| A52 | 106.33950994 | 106.35485078 |
| A53 | 109.65684048 | 109.75541245 |
| A54 | 62.50431090 | 62.44180079 |
| A55 | 79.14840553 | 78.61332450 |
| A56 | 94.38442272 | 93.89685773 |
| A57 | 81.94439843 | 81.33655214 |
| A58 | 122.30128587 | 122.38085758 |
| A59 | 108.69599817 | 108.46009440 |
| A60 | 118.51678659 | 118.96829710 |
| A61 | 112.14562665 | 112.66775833 |
| A62 | 125.70871686 | 125.55440414 |
| A63 | 104.10605874 | 104.05408726 |
| A64 | 115.66124171 | 115.41491837 |
| A65 | 112.85938504 | 112.90944315 |
| A66 | 111.09046713 | 111.04846793 |
| A67 | 109.12827188 | 109.12112805 |
| A68 | 112.50720874 | 112.20795577 |
| A69 | 108.93102542 | 108.90860024 |
| A70 | 109.76379473 | 109.82714850 |
| A71 | 35.55295345 | 35.64055684 |
| A72 | 108.78209604 | 109.15267108 |
| A73 | 35.76405946 | 35.85622083 |
| A74 | 104.88212372 | 105.29541039 |
| D1 | 23.44707166 | 22.86899643 |
| D2 | −143.01621213 | −145.69511584 |
| D3 | 38.42108435 | 38.12739399 |
| D4 | 167.03950592 | 166.71102018 |
| D5 | 11.19248174 | 10.26295325 |
| D6 | −166.27299883 | −167.50854474 |
| D7 | 154.43296848 | 154.87100444 |
| D8 | −24.72393257 | −23.90813959 |
| D9 | 14.87760390 | 14.33966315 |
| D10 | −166.33850331 | −167.39527774 |
| D11 | −178.68215577 | −179.05559597 |
| D12 | 116.12498790 | 117.52135961 |
| D13 | −96.95997467 | −97.21214362 |
| D14 | 10.99862661 | 10.70745397 |
| D15 | 143.71999664 | 143.27578873 |
| D16 | 53.45687997 | 56.95553385 |
| D17 | −13.05739505 | −15.48602055 |
| D18 | 166.77990428 | 164.20618170 |
| D19 | 179.92648481 | 179.96617851 |
| D20 | −0.52972049 | −0.55875352 |
| D21 | −179.66796941 | −179.58911616 |
| D22 | 0.73133137 | 0.86390555 |
| D23 | 179.16217965 | 179.06507149 |
| D24 | 0.01070889 | −0.03915718 |
| D25 | 179.03888636 | 179.05702057 |
| D26 | 48.07112323 | 45.04719980 |
| D27 | 0.74469490 | 0.91641539 |
| D28 | −179.96524676 | −179.74189717 |
| D29 | −179.85477901 | −179.85236519 |
| D30 | −0.39506461 | −0.41589868 |
| D31 | −179.79735693 | −179.79071017 |
| D32 | 0.01649873 | 0.03030576 |
| D33 | 179.92823773 | 179.93269511 |
| D34 | 0.28491317 | 0.27631314 |
| D35 | 179.18887066 | 179.19810092 |
| D36 | −116.75958591 | −117.79096246 |
| D37 | 1.86219749 | 1.56294201 |
| D38 | −115.86020598 | −117.02116664 |
| D39 | −179.58717365 | −179.74490671 |
| D40 | 63.79739074 | 62.83877093 |
| D41 | 178.95201872 | 179.06783274 |
| D42 | −0.28698798 | −0.13593353 |
| D43 | 178.22649816 | 178.07298377 |
| D44 | −148.93368537 | −152.08506467 |
| D45 | 111.42783975 | 113.18142815 |
| D46 | −4.05565899 | −2.40606604 |
| D47 | −179.15665881 | −177.51084506 |
| D48 | −12.57964549 | −12.42277636 |
| D49 | −120.44409694 | −120.19086848 |
| D50 | 98.09202903 | 96.64992442 |
| D51 | 91.15604637 | 91.57000780 |
| D52 | −153.66577605 | −153.20632535 |
| D53 | 135.69337038 | 135.76291529 |
| D54 | 154.19232954 | 153.37488771 |
| D55 | −1.14110681 | −0.79096165 |
| D56 | 4.84906073 | 4.80879280 |
| D57 | −83.86223058 | −85.19930682 |
| D58 | −1.48047919 | −1.48833544 |
| D59 | 161.82475625 | 161.81654714 |
| D60 | 8.46484636 | 8.87965902 |
| D61 | 147.77400083 | 149.20907105 |
| D62 | −178.10763280 | −178.26760363 |
| D63 | 66.16427504 | 65.99502020 |
| D64 | −71.70727202 | −71.00674962 |
| D65 | 50.12782255 | 50.90167682 |
| D66 | 168.95273147 | 169.66688376 |
| D67 | −126.11657682 | −124.36123085 |
| D68 | 54.49143257 | 54.30008219 |
| D69 | −62.36290369 | −62.63339700 |
| D70 | −66.30729976 | −66.82133358 |
| D71 | −118.50510691 | −118.64472010 |
| D72 | −128.64978340 | −128.59930105 |
| D73 | −115.36964313 | −115.74853701 |

**Table A3.**
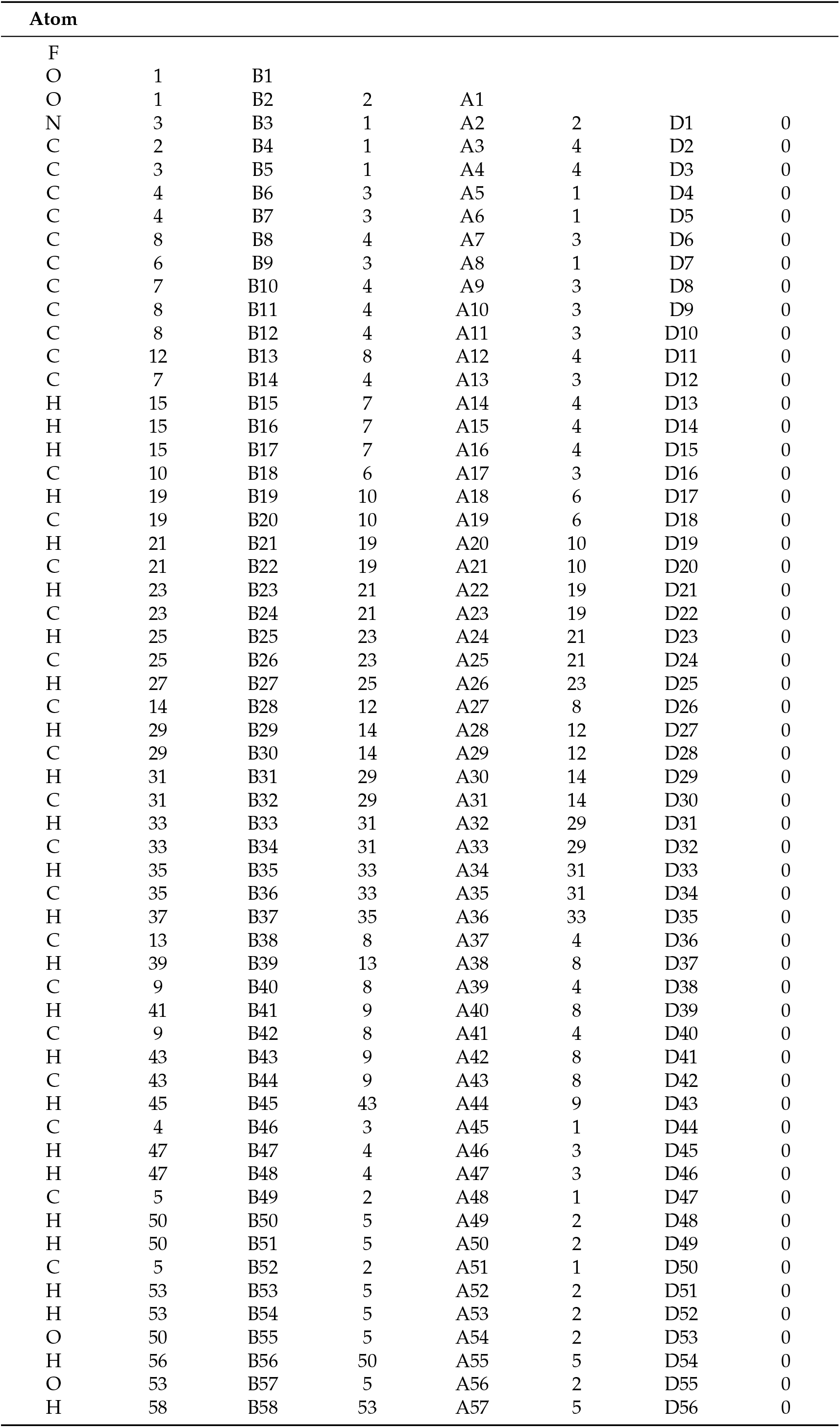
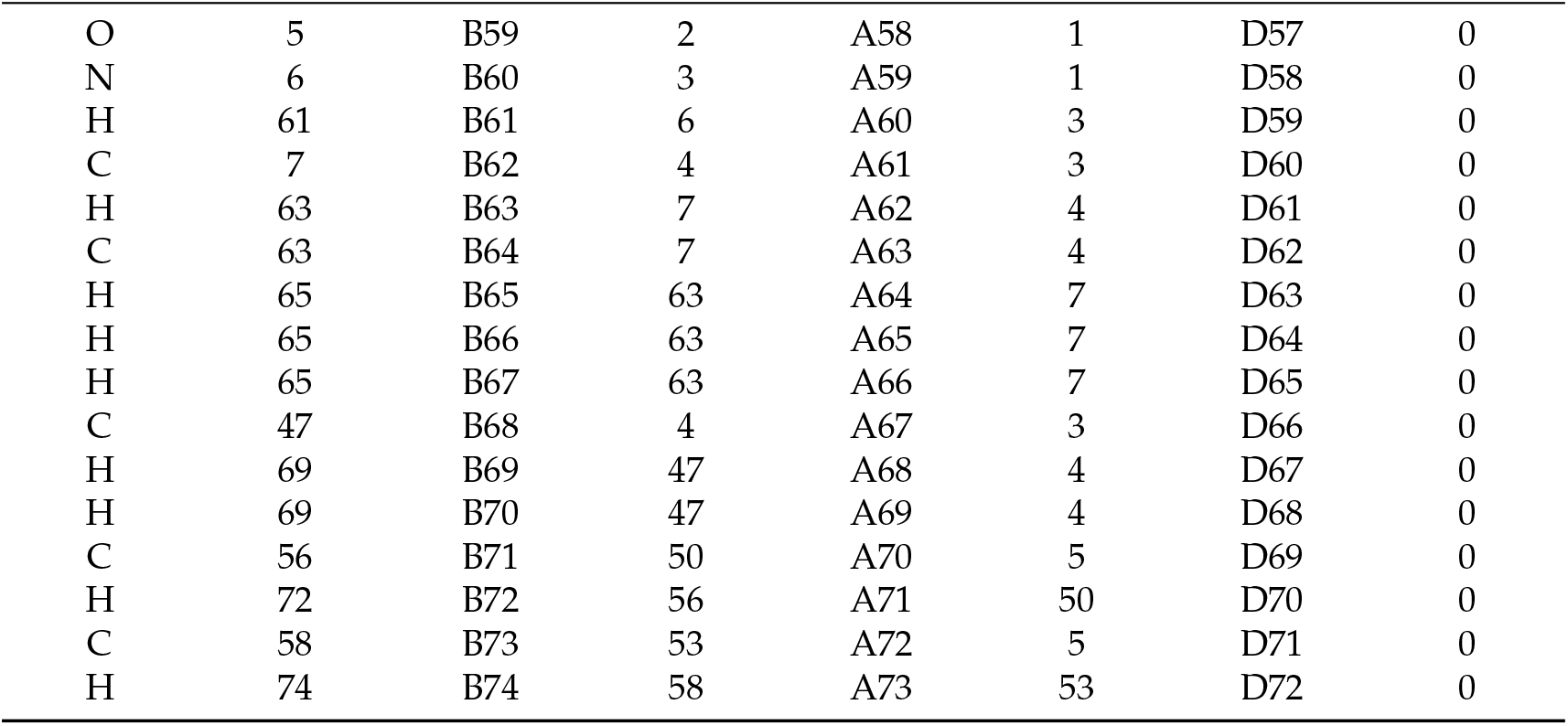
Z−matrix of ATV1H.

| Atom |  |  |  |  |  |  |  |
| --- | --- | --- | --- | --- | --- | --- | --- |
| F |  |  |  |  |  |  |  |
| O | 1 | B1 |  |  |  |  |  |
| O | 1 | B2 | 2 | A1 |  |  |  |
| N | 3 | B3 | 1 | A2 | 2 | D1 | 0 |
| C | 2 | B4 | 1 | A3 | 4 | D2 | 0 |
| C | 3 | B5 | 1 | A4 | 4 | D3 | 0 |
| C | 4 | B6 | 3 | A5 | 1 | D4 | 0 |
| C | 4 | B7 | 3 | A6 | 1 | D5 | 0 |
| C | 8 | B8 | 4 | A7 | 3 | D6 | 0 |
| C | 6 | B9 | 3 | A8 | 1 | D7 | 0 |
| C | 7 | B10 | 4 | A9 | 3 | D8 | 0 |
| C | 8 | B11 | 4 | A10 | 3 | D9 | 0 |
| C | 8 | B12 | 4 | A11 | 3 | D10 | 0 |
| C | 12 | B13 | 8 | A12 | 4 | D11 | 0 |
| C | 7 | B14 | 4 | A13 | 3 | D12 | 0 |
| H | 15 | B15 | 7 | A14 | 4 | D13 | 0 |
| H | 15 | B16 | 7 | A15 | 4 | D14 | 0 |
| H | 15 | B17 | 7 | A16 | 4 | D15 | 0 |
| C | 10 | B18 | 6 | A17 | 3 | D16 | 0 |
| H | 19 | B19 | 10 | A18 | 6 | D17 | 0 |
| C | 19 | B20 | 10 | A19 | 6 | D18 | 0 |
| H | 21 | B21 | 19 | A20 | 10 | D19 | 0 |
| C | 21 | B22 | 19 | A21 | 10 | D20 | 0 |
| H | 23 | B23 | 21 | A22 | 19 | D21 | 0 |
| C | 23 | B24 | 21 | A23 | 19 | D22 | 0 |
| H | 25 | B25 | 23 | A24 | 21 | D23 | 0 |
| C | 25 | B26 | 23 | A25 | 21 | D24 | 0 |
| H | 27 | B27 | 25 | A26 | 23 | D25 | 0 |
| C | 14 | B28 | 12 | A27 | 8 | D26 | 0 |
| H | 29 | B29 | 14 | A28 | 12 | D27 | 0 |
| C | 29 | B30 | 14 | A29 | 12 | D28 | 0 |
| H | 31 | B31 | 29 | A30 | 14 | D29 | 0 |
| C | 31 | B32 | 29 | A31 | 14 | D30 | 0 |
| H | 33 | B33 | 31 | A32 | 29 | D31 | 0 |
| C | 33 | B34 | 31 | A33 | 29 | D32 | 0 |
| H | 35 | B35 | 33 | A34 | 31 | D33 | 0 |
| C | 35 | B36 | 33 | A35 | 31 | D34 | 0 |
| H | 37 | B37 | 35 | A36 | 33 | D35 | 0 |
| C | 13 | B38 | 8 | A37 | 4 | D36 | 0 |
| H | 39 | B39 | 13 | A38 | 8 | D37 | 0 |
| C | 9 | B40 | 8 | A39 | 4 | D38 | 0 |
| H | 41 | B41 | 9 | A40 | 8 | D39 | 0 |
| C | 9 | B42 | 8 | A41 | 4 | D40 | 0 |
| H | 43 | B43 | 9 | A42 | 8 | D41 | 0 |
| C | 43 | B44 | 9 | A43 | 8 | D42 | 0 |
| H | 45 | B45 | 43 | A44 | 9 | D43 | 0 |
| C | 4 | B46 | 3 | A45 | 1 | D44 | 0 |
| H | 47 | B47 | 4 | A46 | 3 | D45 | 0 |
| H | 47 | B48 | 4 | A47 | 3 | D46 | 0 |
| C | 5 | B49 | 2 | A48 | 1 | D47 | 0 |
| H | 50 | B50 | 5 | A49 | 2 | D48 | 0 |
| H | 50 | B51 | 5 | A50 | 2 | D49 | 0 |
| C | 5 | B52 | 2 | A51 | 1 | D50 | 0 |
| H | 53 | B53 | 5 | A52 | 2 | D51 | 0 |
| H | 53 | B54 | 5 | A53 | 2 | D52 | 0 |
| O | 50 | B55 | 5 | A54 | 2 | D53 | 0 |
| H | 56 | B56 | 50 | A55 | 5 | D54 | 0 |
| O | 53 | B57 | 5 | A56 | 2 | D55 | 0 |
| H | 58 | B58 | 53 | A57 | 5 | D56 | 0 |
| O | 5 | B59 | 2 | A58 | 1 | D57 | 0 |
| N | 6 | B60 | 3 | A59 | 1 | D58 | 0 |
| H | 61 | B61 | 6 | A60 | 3 | D59 | 0 |
| C | 7 | B62 | 4 | A61 | 3 | D60 | 0 |
| H | 63 | B63 | 7 | A62 | 4 | D61 | 0 |
| C | 63 | B64 | 7 | A63 | 4 | D62 | 0 |
| H | 65 | B65 | 63 | A64 | 7 | D63 | 0 |
| H | 65 | B66 | 63 | A65 | 7 | D64 | 0 |
| H | 65 | B67 | 63 | A66 | 7 | D65 | 0 |
| C | 47 | B68 | 4 | A67 | 3 | D66 | 0 |
| H | 69 | B69 | 47 | A68 | 4 | D67 | 0 |
| H | 69 | B70 | 47 | A69 | 4 | D68 | 0 |
| C | 56 | B71 | 50 | A70 | 5 | D69 | 0 |
| H | 72 | B72 | 56 | A71 | 50 | D70 | 0 |
| C | 58 | B73 | 53 | A72 | 5 | D71 | 0 |
| H | 74 | B74 | 58 | A73 | 53 | D72 | 0 |

**Table A4.**
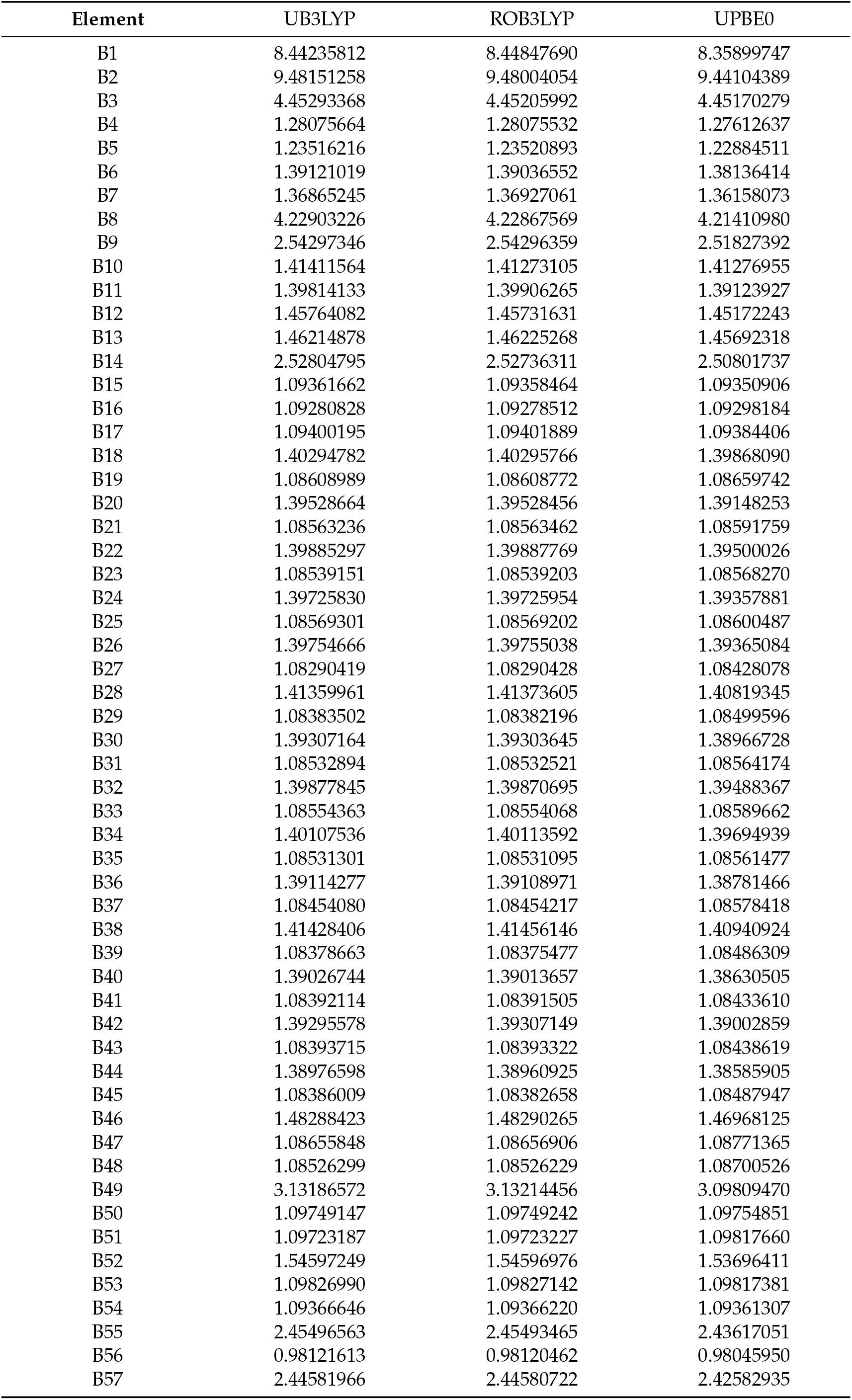
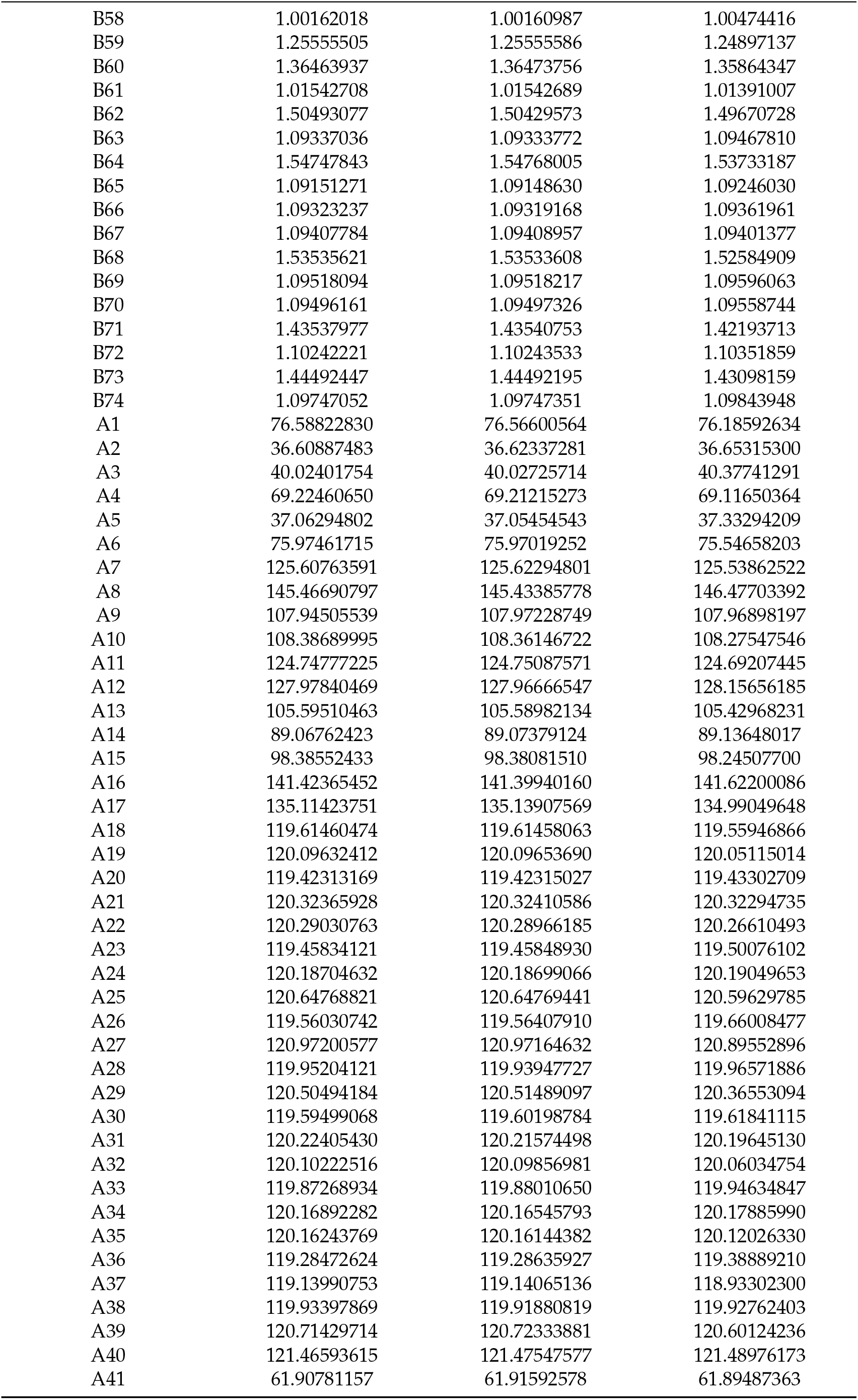
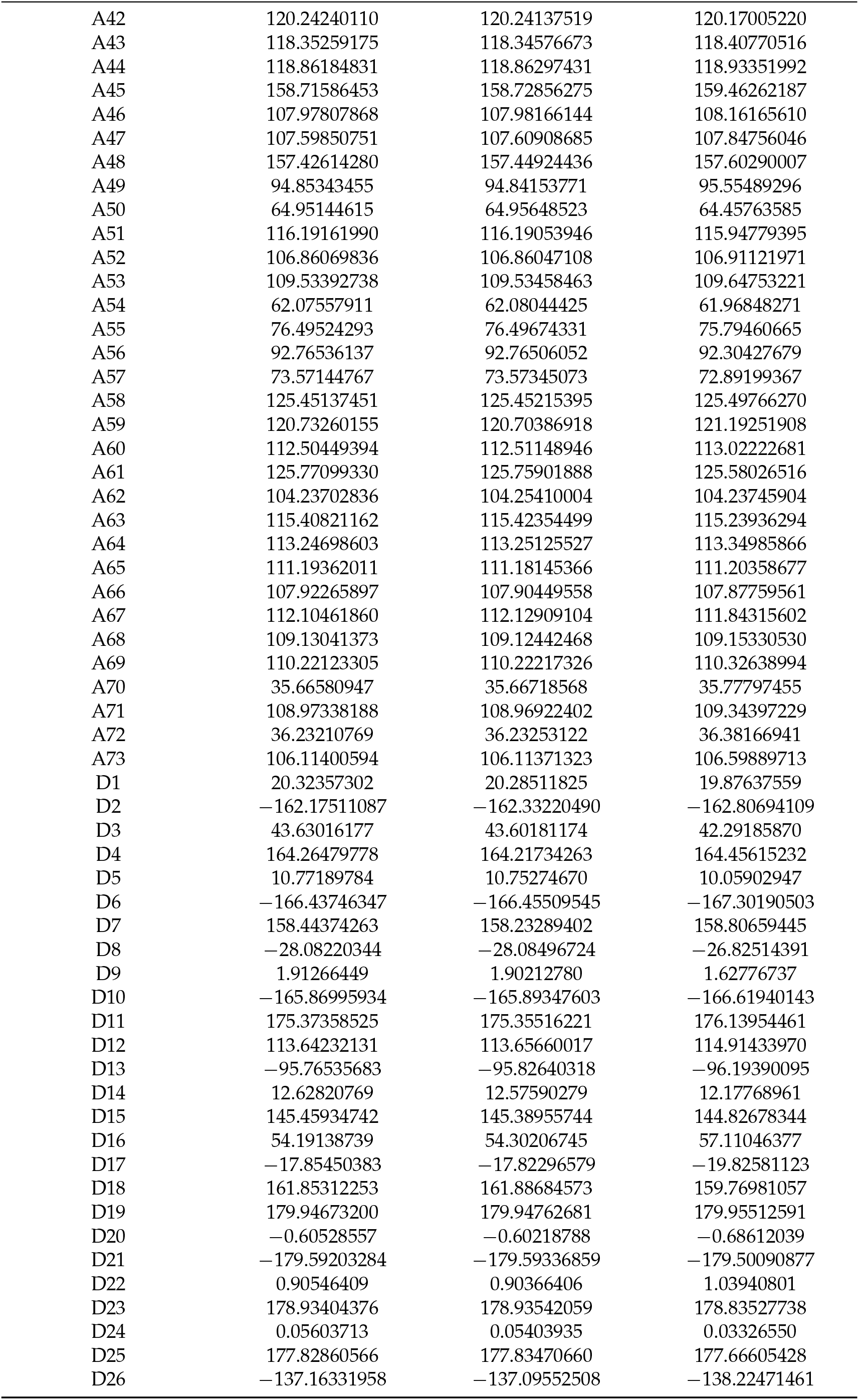
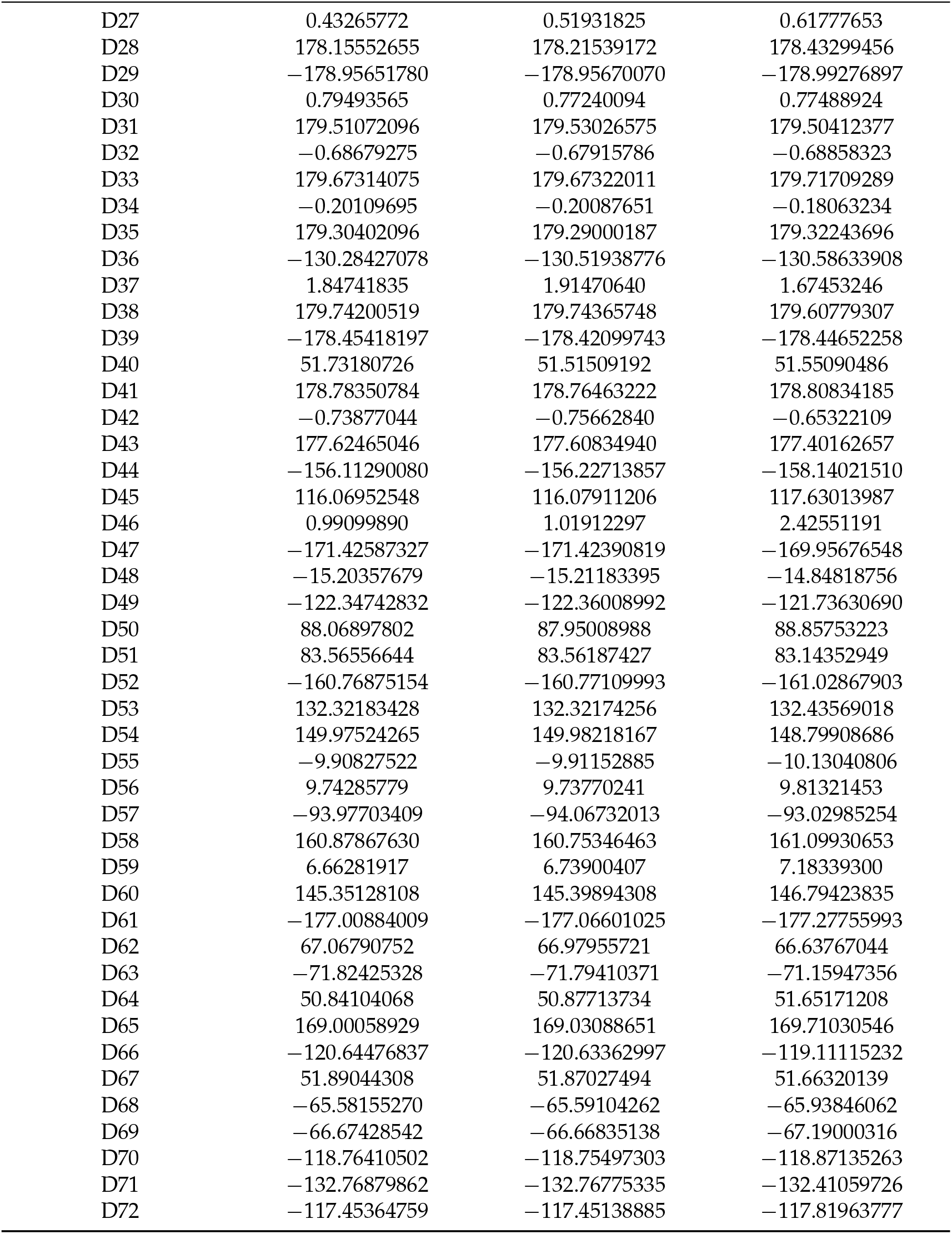
Elements of the Z−matrix of ATV1H in methanol optimized as indicated below using 6-31+G/d,p) basis sets.

